# Population-associated molecular variation in histologically normal breast tissue is context-dependent and associated with distinct transcriptional states

**DOI:** 10.64898/2026.06.16.732684

**Authors:** William Drew Hulsy, Karen Salazar, Dimitra Chalkia, Yonny Chavez, Yuchen Zhao, Georgia Halkia, Olga V. Razorenova, Nikolas Nikolaidis

## Abstract

Population-associated molecular variation in breast tissue may contribute to differences in tissue biology and disease susceptibility, yet the extent to which such variation is shaped by underlying tissue states remains unclear. Here, we performed RNA-seq and lipidomic profiling of histologically normal breast tissue samples from African American (AA) and Caucasian White (CW) individuals, followed by conceptual integration of the resulting transcriptomic and lipidomic patterns. Unsupervised analysis revealed two distinct baseline transcriptional states (G1 and G2) that defined the primary axis of molecular variation across the cohort and corresponded to epithelial-enriched (G1) and vascular-enriched (G2) tissue contexts as determined by cell-type deconvolution. Global comparisons between AA and CW samples showed minimal transcriptomic differences, with only a single gene reaching significance after multiple testing correction. However, when stratified by baseline tissue state, 191 genes were differentially expressed within G1, with coordinated upregulation of extracellular matrix organization and proliferative/cytoskeletal processes in AA samples. These patterns were consistently supported across multiple enrichment approaches. No comparable population-associated differences were observed within G2. Lipidomic analyses showed partial but non-significant trends consistent with transcriptomic structure, suggesting that lipid variation provides complementary but limited support for baseline molecular differences, likely reflecting constraints of bulk tissue composition. Together, these findings suggest that population-associated molecular differences in normal breast tissue are context-dependent and emerge within specific baseline transcriptional states, where distinct biological programs can coexist and be differentially modulated. These findings highlight the importance of tissue heterogeneity in shaping molecular variation and its potential relevance to disease-associated tissue states.

## 1. Introduction

Human populations exhibit measurable molecular variation across individuals, reflecting interactions among genetic ancestry, environmental exposure, socioeconomic context, and physiological history [1–3]. African American women experience disproportionately higher breast cancer mortality compared to Caucasian White women, a disparity that persists after accounting for stage at diagnosis and treatment access, suggesting that biological factors may contribute alongside clinical and socioeconomic determinants [1–4]. Differences observed across population groups may therefore arise from a combination of biological and lived factors rather than any single determinant [2]. While population differences in tumor biology and treatment response have received considerable attention [1–3], little is known about how baseline molecular programs differ in histologically normal tissues across population groups [5–7]. Characterizing such variation is essential for defining the range of normal biological states and for understanding how tissue context may differ prior to disease onset [6,8].

Normal breast tissue represents a structurally and cellularly complex system composed of epithelial, stromal, immune, and adipose compartments [1,8]. This heterogeneity contributes to continuous remodeling across the lifespan and in response to hormonal and metabolic cues [1,9]. Despite its biological relevance, baseline molecular variation in non-tumor breast tissue remains incompletely characterized, particularly across diverse population groups [5–7]. In bulk tissue analyses, distinguishing intrinsic transcriptional programs from differences in cellular composition or tissue architecture is especially important, as variation in cellular composition and tissue architecture can strongly influence observed molecular patterns [1,8].

Integrated multi-omic approaches provide a framework for interrogating baseline tissue states across multiple molecular layers [10–12]. Transcriptomic profiling can identify coordinated biological programs such as extracellular matrix organization, cytoskeletal dynamics, and proliferative activity [1,8], while lipidomic profiling offers complementary insight into metabolic context and tissue composition, as shown in breast tissue lipidomics studies [10,13]. In breast tissue specifically, which is adipose-rich, metabolically active, and hormonally responsive, lipid composition reflects both the cellular milieu and systemic physiological state, making it a particularly informative complement to transcriptomic data [8,10,13]. In this study, we use conceptual integration to refer to the independent analysis of each omics layer followed by biological comparison of the resulting transcriptomic and lipidomic patterns, rather than formal feature-level statistical integration across platforms [8,11,12].

Here, we present a pilot multi-omic analysis of histologically normal breast tissue samples from African American and Caucasian White women, using RNA-seq and lipidomic profiling followed by conceptual integration of the resulting molecular patterns. We specifically focused on these two population groups given the well-documented breast cancer outcome disparities between them, with the goal of determining whether baseline differences in normal tissue molecular programs, prior to any disease process, can be detected and contextualized within underlying tissue states. Given the emerging role of lipid metabolism as a driver of metastasis and poor survival in triple-negative breast cancer [14–16], we included lipidomic profiling to complement transcriptional analysis and explore whether baseline lipid programs differ between population groups in normal breast tissue. With a modest cohort size and inherent heterogeneity of bulk tissue, the primary aim was to identify coherent transcriptional and metabolic patterns rather than to establish causal mechanisms.

## 2. Results

### 2.1. Cohort Structure and Analytical Overview

#### 2.1.1 Cohort composition and sample matching

The study cohort consisted of 30 histologically normal breast tissue samples obtained from individuals self-identified as African American (AA; n = 14) or Caucasian White (CW; n = 16) (Table 1 and Methods). Samples were selected under demographic matching criteria, with particular attention to age and body mass index (BMI), to reduce major confounding effects (Figure 1). Due to limitations in sample availability, matching required selective inclusion of lower-BMI individuals in the AA group to achieve overlap between groups. As such, the analyzed cohort represents a demographically constrained subset rather than a fully representative population sample.

**Figure 1.**
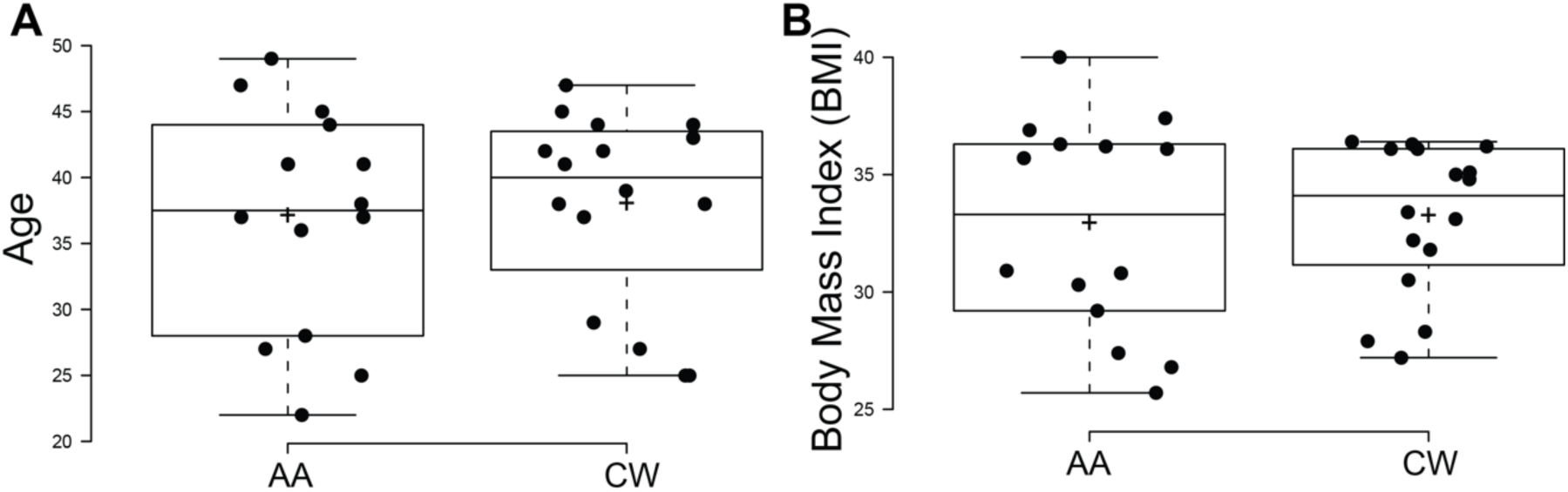
Cohort demographic matching. (A) Age distribution and (B) body mass index (BMI) distribution of samples by population group (AA, African American; CW, Caucasian White). Samples included in the study were selected to achieve comparable distributions of age and BMI between population groups to reduce major demographic confounding. Box plots indicate the median and interquartile range, with the mean shown as a cross and individual samples displayed as points (AA, n = 14; CW, n = 16). Statistical comparisons were performed using a two-sided Mann–Whitney U test. No statistically significant differences were observed between groups (Age: p = 0.766; BMI: p = 0.198), indicating comparable demographic distributions across the analyzed cohort.

**Table 1.** Cohort metadata for normal human breast tissue samples included in the multi-omic analysis. Population groups were defined based on self-identified population (AA, African American; CW, Caucasian White). Transcriptomic clustering assigned a subset of samples into two baseline tissue states (G1 and G2) based on unsupervised analysis. Samples not assigned to either group are indicated as NA. Age and body mass index (BMI) were used for cohort matching to minimize major demographic confounding

| Age | BMI | Sample Label | Population | Group |
| --- | --- | --- | --- | --- |
| 42 | 27.9 | CW1 | CW | G1 |
| 37 | 34.8 | CW10 | CW | G1 |
| 38 | 35.1 | CW11 | CW | G1 |
| 44 | 31.8 | CW6 | CW | G1 |
| 41 | 36.1 | CW7 | CW | G1 |
| 27 | 27.2 | CW9 | CW | G1 |
| 28 | 36.2 | AA1 | AA | G1 |
| 25 | 30.9 | AA4 | AA | G1 |
| 44 | 29.2 | AA5 | AA | G1 |
| 41 | 36.1 | AA9 | AA | G1 |
| 45 | 36.3 | CW12 | CW | G2 |
| 43 | 33.1 | CW2 | CW | G2 |
| 29 | 36.1 | CW3 | CW | G2 |
| 25 | 35.0 | CW4 | CW | G2 |
| 38 | 36.4 | CW8 | CW | G2 |
| 49 | 30.3 | AA10 | AA | G2 |
| 27 | 27.4 | AA11 | AA | G2 |
| 41 | 35.7 | AA2 | AA | G2 |
| 38 | 36.3 | AA3 | AA | G2 |
| 47 | 26.8 | AA6 | AA | G2 |
| 37 | 36.9 | AA7 | AA | G2 |
| 37 | 37.4 | AA8 | AA | G2 |
| 39 | 33.4 | CW13 | CW | NA |
| 42 | 36.2 | CW14 | CW | NA |
| 47 | 28.3 | CW15 | CW | NA |
| 25 | 30.5 | CW16 | CW | NA |
| 44 | 32.2 | CW5 | CW | NA |
| 45 | 30.8 | AA12 | AA | NA |
| 36 | 25.7 | AA13 | AA | NA |
| 22 | 40.0 | AA14 | AA | NA |

All 30 samples underwent lipidomic profiling under uniform extraction and analytical conditions (see Methods). For transcriptomic profiling, one sample failed library preparation due to RNA degradation and seven additional samples were excluded based on high duplication rates identified during quality control (see Methods and Supplementary Table S1); the remaining 22 samples were retained for RNA-seq analysis. Of these, unsupervised transcriptomic clustering assigned 10 samples to Group 1 (G1; AA = 4, CW = 6) and 12 to Group 2 (G2; AA = 7, CW = 5). RNA quality metrics and sequencing parameters did not indicate systematic technical separation among retained samples (see Methods and Supplementary Figure S1). Samples excluded from transcriptomic analysis are indicated as NA in Table 1 and were retained for lipidomic comparisons only.

#### 2.1.2 Multi-omic analytical framework

Transcriptomic data were processed using the nf-core/rnaseq pipeline (v3.14.0), with differential expression analysis performed using DESeq2 and functional enrichment assessed across multiple platforms including Gene Ontology over-representation analysis (Enrichr), Gene Set Enrichment Analysis (GSEA using fgsea against MSigDB collections), topGO (weight01 algorithm, Fisher’s exact test), and STRING network analysis (v12.0, interaction confidence ≥ 0.7). Lipidomic data were analyzed on mTIC-normalized intensities from 252 identified lipid species using both univariate statistical comparisons (unpaired t-test, Benjamini–Hochberg FDR correction) and multivariate approaches, including principal component analysis (PCA) and sparse partial least squares discriminant analysis (sPLS-DA). Integration across modalities was performed at the level of biological program convergence rather than direct gene–lipid pairing.

Given the pilot nature of the study and the modest sample size, the analytical strategy prioritized internal biological consistency across enrichment platforms over reliance on single statistical contrasts. Findings were interpreted conservatively, with emphasis on reproducible pathway-level patterns rather than isolated feature-level differences.

#### 2.1.3 Unsupervised structure of the transcriptomic data

Unsupervised PCA of transcriptomic profiles revealed separation of samples into two primary clusters, hereafter referred to as Group 1 (G1) and Group 2 (G2), along PC1, which accounted for 66% of total transcriptomic variance (Figure 2A). This clustering was reproduced by t-SNE analysis and was not explained by population group assignment, as both AA and CW samples were distributed across each cluster (Figure 2B and C). The G1/G2 separation was also not consistently associated with available metadata variables including age, BMI, or storage categories (Supplementary Figure S1), suggesting the presence of intrinsic baseline transcriptional heterogeneity within the cohort.

**Figure 2.**
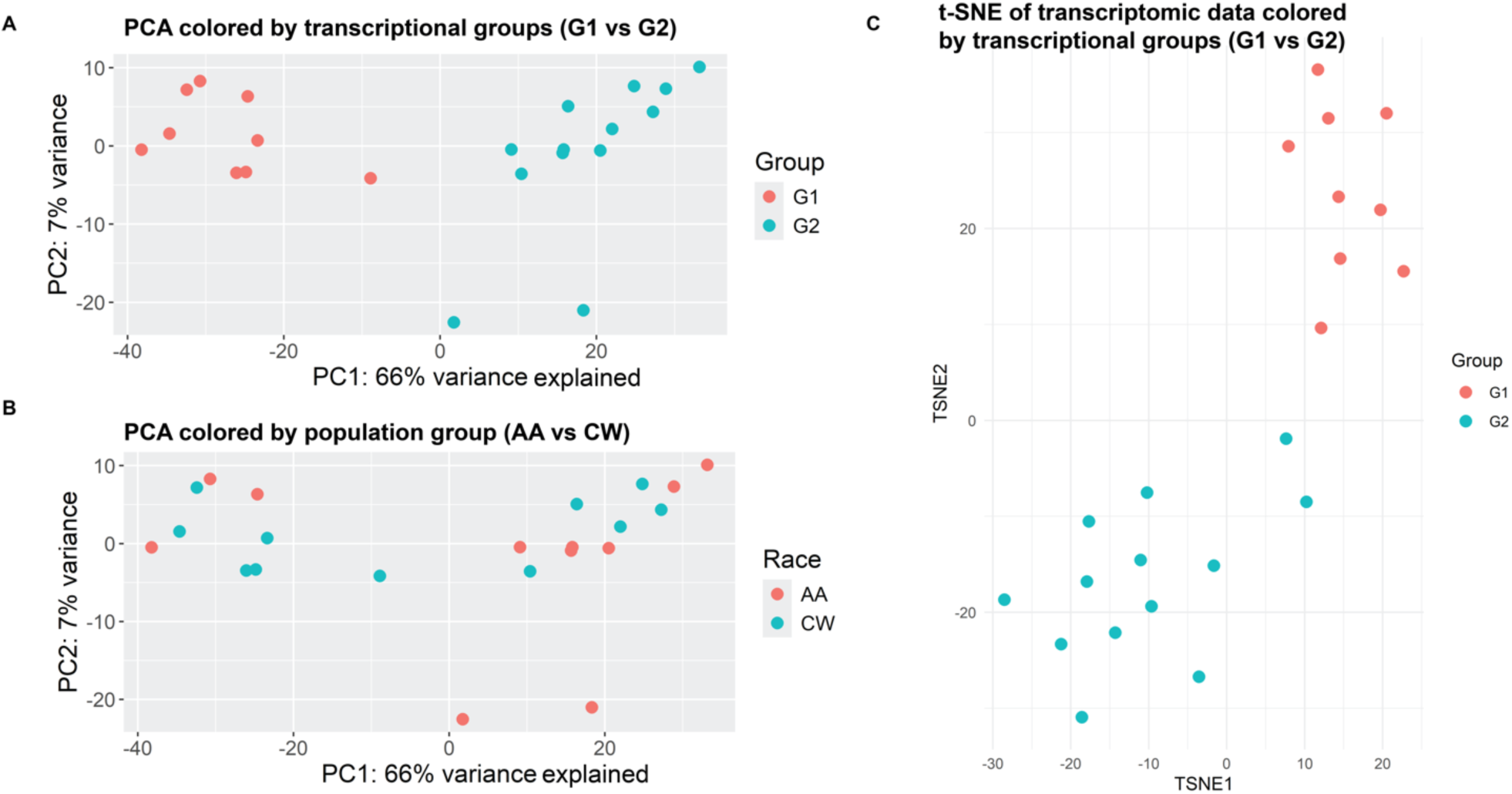
Unsupervised structure of the transcriptomic dataset (n=22). (A) Principal component analysis (PCA) of normalized transcriptomic profiles showing separation of samples into two clusters designated Group 1 (G1) and Group 2 (G2). PC1 accounts for 66% of total variance and drives the primary separation between groups. (B) The same PCA coordinates as in (A), colored by population group (AA, African American; CW, Caucasian White), demonstrating that both population groups are represented within each cluster and therefore the observed separation is not driven by population assignment. (C) t-distributed stochastic neighbor embedding (t-SNE with perplexity = 7) of the same dataset reveals a clustering pattern consistent with the PCA-defined G1 and G2 groups.

Importantly, the emergence of G1 and G2 was data-driven and not defined by population group assignment. This unsupervised structure provided a framework for subsequent within-group and between-group analyses, allowing assessment of whether population-associated differences were globally distributed or context-dependent within specific baseline tissue states.

#### 2.1.4 Overview of comparison strategy

Based on the observed unsupervised clustering, subsequent analyses were structured along two axes:

1. Comparisons between G1 and G2 to characterize intrinsic baseline transcriptional states.
2. Comparisons between population groups both across the full cohort and within specific baseline transcriptional states.

This hierarchical approach enabled evaluation of whether population-associated molecular differences were globally present or confined to specific baseline transcriptional contexts.

### 2.2. Two Baseline Tissue States Dominate Transcriptomic Variation

#### 2.2.1 Transcriptomic and compositional differences between G1 and G2

The G1/G2 transcriptomic separation established by PCA (Figure 2) was further characterized by differential expression analysis. Differential expression analysis between G1 and G2 (|log₂FC (LFC)| ≥ 1, adjusted p-value ≤ 0.05) identified 4,139 significant genes across 25,560 tested, of which 2,503 and 1,636 were up- and down-regulated, respectively, in G1 (Figure 3 and Supplementary Tables S1 and S2). This large number of differentially expressed genes is consistent with the magnitude of separation observed along PC1 and suggests that G1 and G2 represent fundamentally distinct transcriptional states.

**Figure 3.**
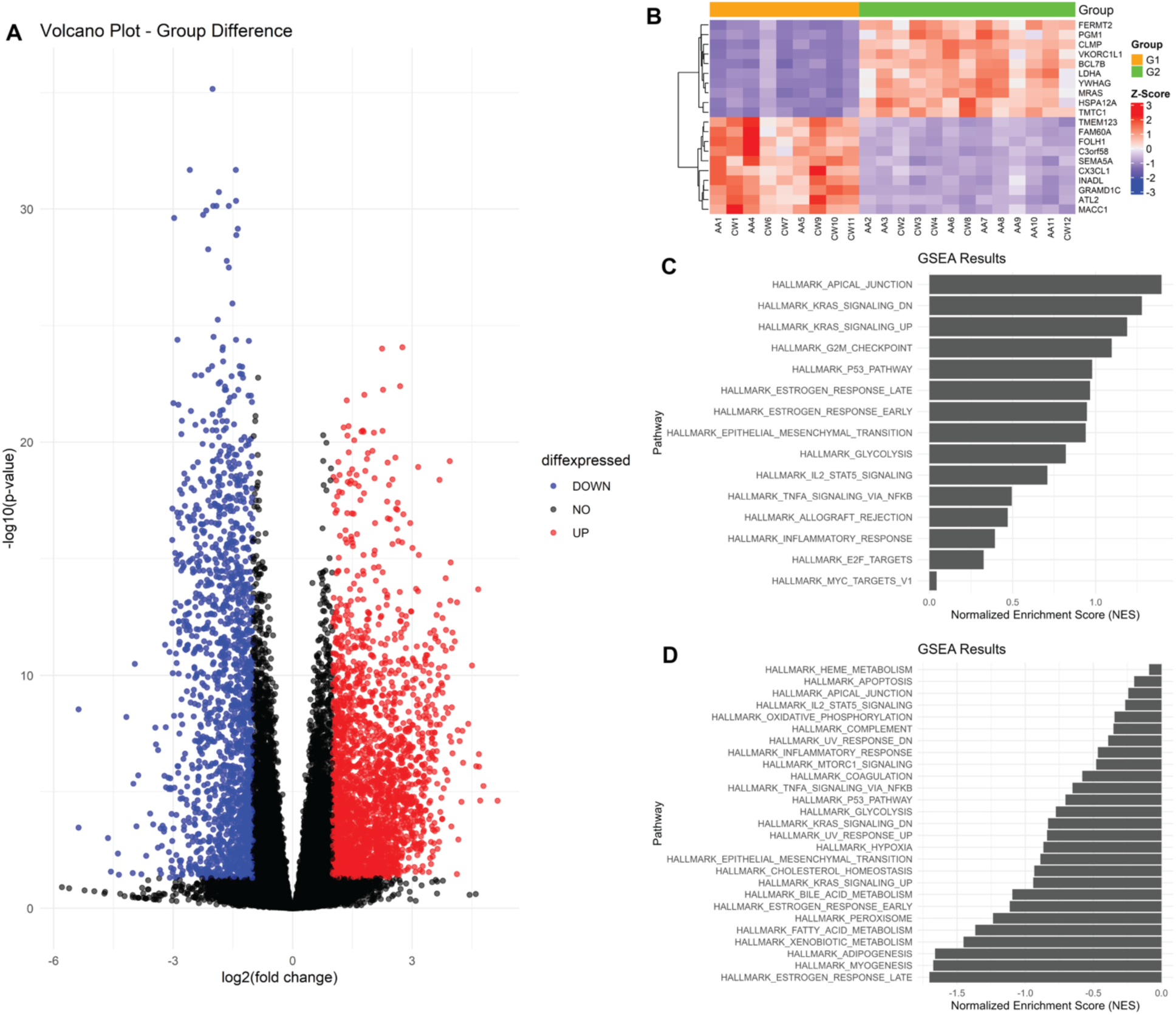
Differential gene expression and pathway enrichment between G1 and G2. (A) Volcano plot showing differentially expressed genes (DEGs) between Group 1 (G1) and Group 2 (G2). Genes are colored based on differential expression status (upregulated, downregulated, or not significant). Significance thresholds were defined as |LFC| ≥ 1 and adjusted p-value < 0.05. (B) Heatmap of the top 10 most significantly up- and down-regulated genes between groups. Genes were ranked by adjusted p-value and displayed with hierarchical clustering of samples. Gene expression values are scaled as Z-scores across samples. Columns represent individual samples and are annotated by group (G1 and G2). (C) Gene set enrichment analysis (GSEA) of Hallmark pathways enriched in G1 relative to G2, shown by normalized enrichment score (NES). (D) GSEA of Hallmark pathways enriched in G2 relative to G1, shown by normalized enrichment score (NES). Bars extend leftward reflecting negative NES values.

GSEA of Hallmark pathways (Supplementary Figure S2 and Supplementary Table S2) identified 15 pathways positively associated with G1 relative to G2, including epithelial junction, proliferative, signaling, and estrogen-response programs (Figure 3C). Pathways enriched in G2 included metabolic, adipogenic, myogenic, and oxidative phosphorylation programs (Figure 3D). Notably, several pathway names appeared in both directions; inspection of the contributing gene sets confirmed that these shared pathway names reflect entirely non-overlapping gene subsets in each direction, indicating that G1 and G2 engage distinct regulatory programs within the same pathway structures. The directional pattern (G1 showing epithelial and proliferative programs and G2 showing metabolic and vascular-associated programs) suggested that the two transcriptional states may reflect underlying differences in cellular composition (Supplementary Table S2).

To evaluate the pathway enrichment results, cell-type deconvolution was performed using MuSiC with a normal human mammary gland single-cell reference [17]. Individual cell types were aggregated into four broad categories: epithelial (luminal epithelial, basal-myoepithelial, and secretory precursor cells), vascular (blood vessel and lymphatic endothelial cells), stromal (fibroblasts and related mesenchymal cells), and immune (leukocytes). G1 samples showed significantly higher estimated epithelial proportions than G2 (median 80.2% vs 60.9%; Wilcoxon rank-sum test, FDR-adjusted p = 0.0003), while G2 samples showed significantly higher estimated vascular proportions (median 36.8% vs 17.6%; adjusted p-value = 0.0003) (Supplementary Table S3). Stromal proportion was higher in G2, yet their difference was not significant. The immune proportion did not differ between the two groups. These compositional differences are consistent with the gene-level enrichment patterns observed in GSEA (Figure 3C–D), linking pathway-level signatures to underlying cellular composition. Because the reference atlas did not contain an annotated adipocyte population, these estimates should be interpreted as relative proportions among the represented epithelial, vascular, stromal, and immune compartments rather than as a complete representation of breast tissue cellular composition.

To determine whether a composition-independent transcriptional signal persisted beyond this compositional difference, differential expression analysis was repeated with estimated epithelial cell-type proportions included as a continuous covariate (design: ∼ epithelial_scaled + group). After adjustment, the expression of 1,268 genes differed significantly between G1 and G2 (adjusted p-value ≤ 0.05, |LFC| ≥ 1; 867 higher in G1, 401 higher in G2), representing a composition-independent transcriptional core. GO Biological Process enrichment analysis using both Enrichr and STRING identified convergent signals across both platforms: G1-enriched genes were consistently associated with ribosomal biogenesis and translational programs (cytoplasmic translation, Enrichr adjusted p-value = 3.52×10⁻⁴⁹, OR = 31.9; STRING p = 3.52×10⁻⁴²), while G2-enriched genes showed concordant enrichment for cell migration, endothelial remodeling, and blood vessel morphogenesis (regulation of focal adhesion assembly, Enrichr adjusted p-value = 1.78×10⁻⁴; blood vessel morphogenesis, STRING p = 2.3×10⁻⁵) (Supplementary Table S4). Together, these findings indicate that G1 and G2 differ at both the level of cellular composition and composition-independent transcriptional programs, with the epithelial-enriched G1 state characterized by high translational output and the vascular-enriched G2 state by active endothelial and migratory programs.

#### 2.2.3 Summary of baseline transcriptional structure

The transcriptomic analyses identified two baseline tissue states, G1 and G2, that represented the dominant axis of molecular variation within the RNA-seq cohort. These states were not attributable to population group, sample storage, or the available demographic variables. Cell-type deconvolution indicated that G1 corresponded to a predominantly epithelial-enriched context, whereas G2 was relatively vascular-enriched. The persistence of substantial differential expression after adjustment for estimated epithelial proportion further indicated that the separation reflected both cellular composition and composition-independent transcriptional programs.

### 2.3. Global Transcriptomic Comparisons between Population Groups

To assess whether molecular differences were present between population groups across the full RNA-seq cohort (n=22), transcriptomic profiles were compared between samples from the two self-identified racial groups without stratification by G1/G2 membership (Figure 2). Principal component analysis of the complete transcriptomic dataset revealed that samples from both population groups were interspersed across the principal component space, indicating that population group assignment alone did not account for the dominant axes of transcriptional variation (Figure 2B).

Differential expression analysis across the full cohort (DESeq2; FDR ≤ 0.1, |LFC| ≥ 1) identified a single gene meeting both thresholds: UNC13C (LFC = −2.60, adjusted p = 9.3 × 10⁻⁵), which showed lower expression in AA samples (Figure 4 and Supplementary Table S5). However, UNC13C showed very low baseline expression across the cohort (detectable above 30 raw counts in only 2 of 22 samples), and this result should therefore be interpreted with caution pending validation in a larger cohort. Given the limited number of significant genes at the standard FDR threshold (≤ 0.05), a relaxed threshold of FDR ≤ 0.1 was applied to capture additional candidate genes for exploratory interpretation. This analysis yielded additional exploratory genes including PDIA3, MIR29B2CHG, ADAMTS9-AS2, LINC-PINT, NR3C2, HLA-F, GPX8, SCN7A, ZNF540, CHIT1, UGDH, CKAP5, LINC01359, and ENSG00000161570, several of which are linked to immune regulation, ion transport, or extracellular matrix remodeling (Figure 4). Functional enrichment analysis of this limited gene set yielded no robust pathway-level signal. No coherent large-scale transcriptional program distinguishing population groups across the entire dataset was evident.

**Figure 4.**
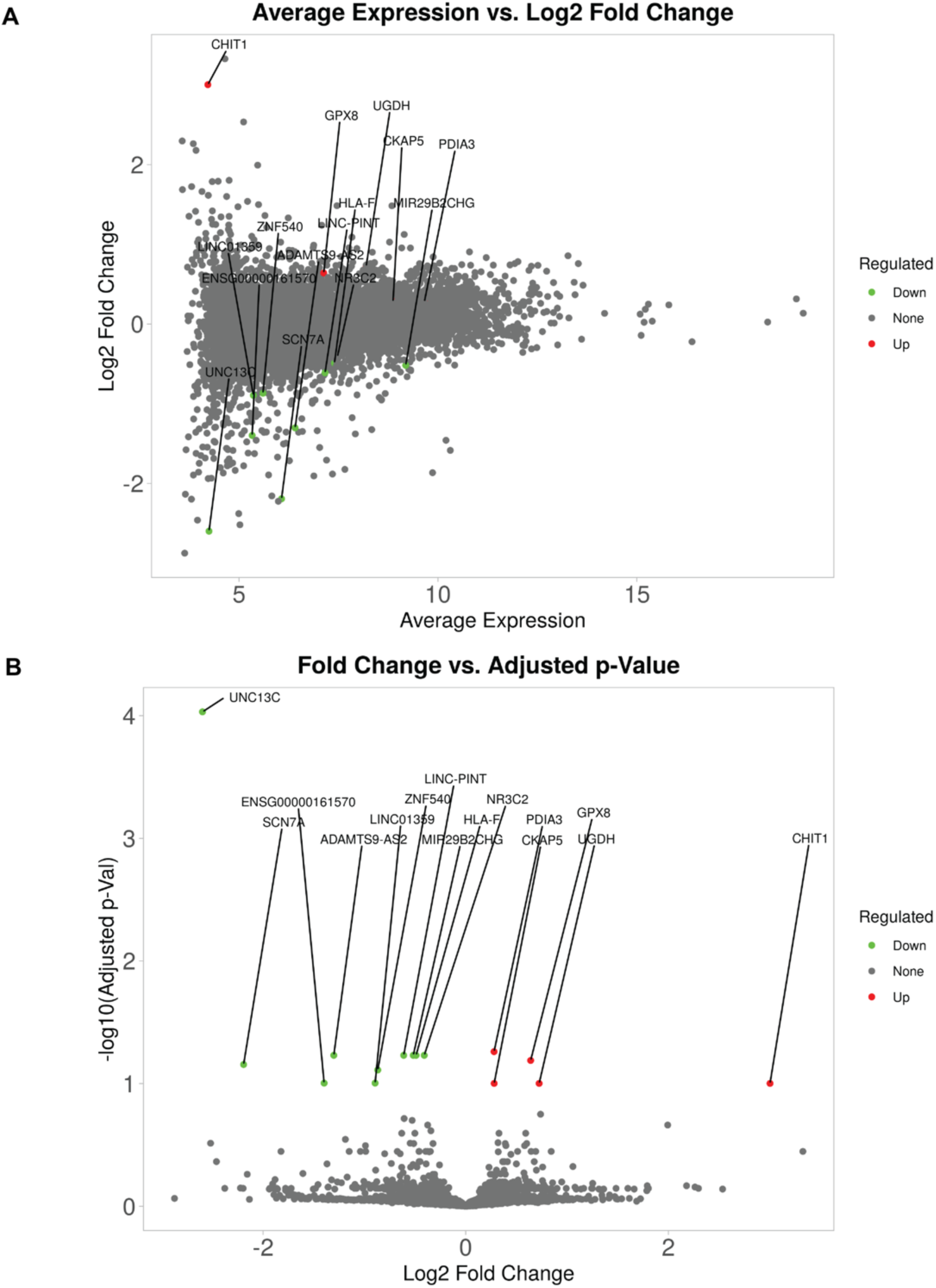
Global transcriptomic comparison between population groups (AA vs CW). (A) MA plot showing LFC versus average expression for all genes comparing African American (AA) and Caucasian White (CW) samples across the full cohort. Genes are colored based on differential expression status using thresholds of adjusted *p*-value ≤ 0.1 and |LFC| ≥ 1. Genes meeting the relaxed threshold (adjusted p-value ≤ 0.1) are labeled and colored; red indicates higher expression in AA, green indicates lower expression in AA. (B) Volcano plot showing LFC versus −log10(adjusted *p*-value) for all genes. Only a single gene, UNC13C, meets both statistical and fold-change thresholds after multiple testing correction. Additional genes labeled represent those with adjusted *p*-value ≤ 0.1 but not necessarily meeting the fold-change cutoff, including PDIA3, MIR29B2CHG, ADAMTS9-AS2, LINC-PINT, NR3C2, HLA-F, GPX8, SCN7A, ZNF540, CHIT1, UGDH, CKAP5, LINC01359, and ENSG00000161570.

Together, these results indicate that population group assignment did not define a dominant axis of transcriptomic variation across the full cohort. Population-associated transcriptional differences, if present, were therefore modest or obscured when samples representing distinct baseline tissue states were analyzed together.

### 2.4. Population-Associated Transcriptomic Differences within Baseline Groups

#### 2.4.1 Transcriptomic differences within G1 and G2

Given the limited global separation between population groups across the full cohort, and the presence of distinct baseline transcriptomic states (G1 and G2), transcriptomic comparisons were next performed within each group (G1, G2). Differential expression analysis within G1 (DESeq2; |LFC| ≥ 1, adjusted p-value ≤ 0.05) identified 191 significantly differentially expressed genes between AA and CW samples (Figure 5 and Supplementary Table S6). Volcano plot analysis confirmed that this within-G1 signal was markedly stronger than that observed in the full cohort comparison, with many genes reaching high statistical confidence (−log₁₀ adjusted p-value > 10) and moderate fold changes (LFC between ±1–3) (Figure 5). In contrast, the equivalent comparison within G2 identified no genes meeting significance thresholds, indicating that population-associated transcriptomic differences were specific to the G1 baseline state (Figure 5B).

**Figure 5.**
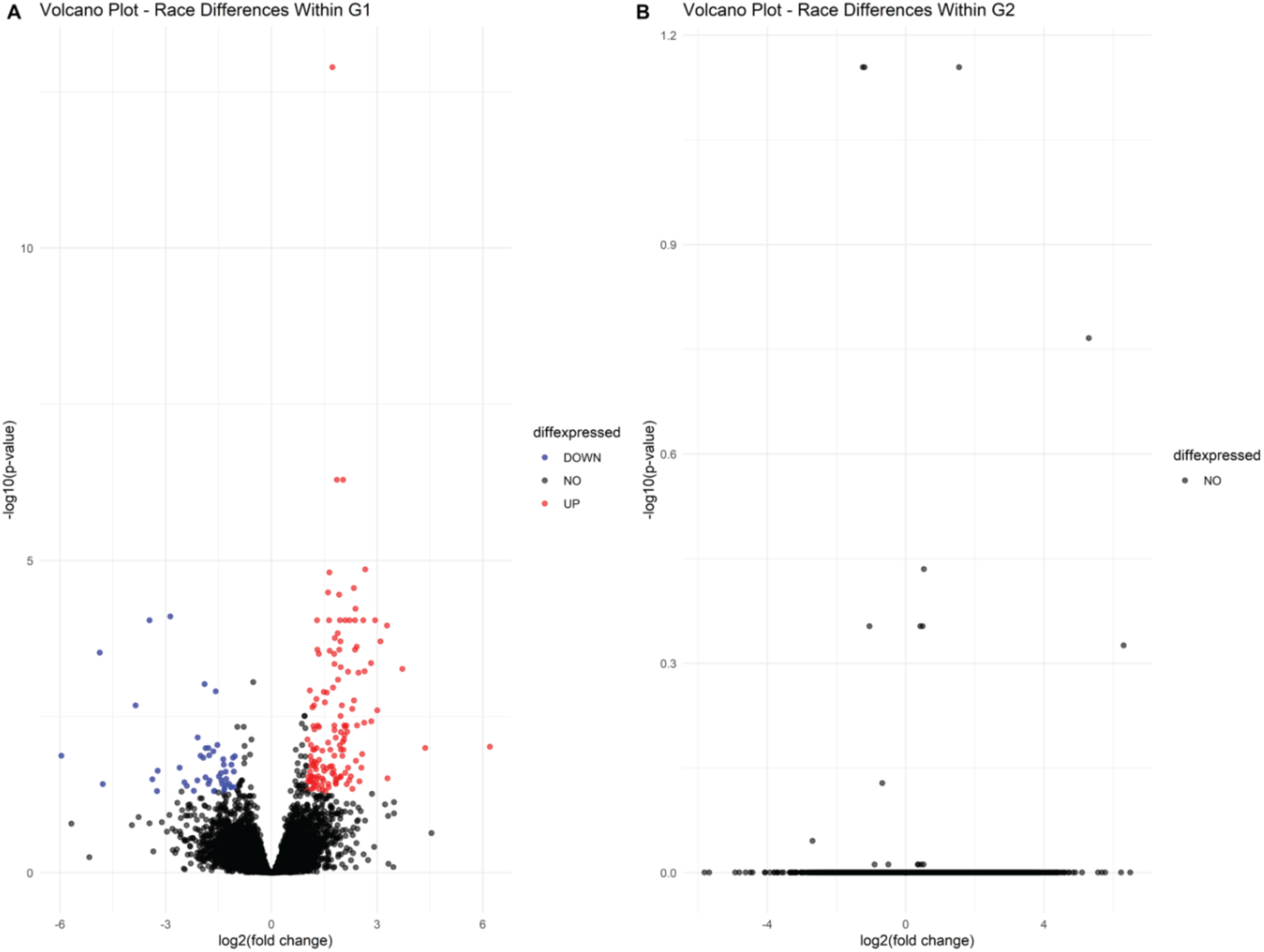
Transcriptomic differences between population groups within G1 and G2. (A) Volcano plot showing differential gene expression between African American (AA) and Caucasian White (CW) samples within Group 1 (G1). Genes are colored based on differential expression status using thresholds of adjusted *p*-value ≤ 0.05 and |LFC| ≥ 1. A substantial number of genes are significantly differentially expressed (191 DEGs), with many showing high statistical significance and moderate fold changes. (B) Volcano plot showing differential gene expression between AA and CW samples within Group 2 (G2) using the same thresholds. In contrast to G1, no genes meet significance criteria in G2.

To assess whether the transcriptional differences could be attributed to differences in cellular composition between AA and CW samples within G1, estimated cell-type proportions derived from MuSiC deconvolution were compared between groups. No significant differences were observed in any cell-type category (all FDR-adjusted p > 0.81; Supplementary Table S3), and median proportions were nearly identical between groups (epithelial: AA 80.0% vs CW 80.2%; vascular: AA 16.2% vs CW 18.8%). These findings indicate that the population-associated transcriptional differences within G1 are unlikely to be explained by differences in the broad cellular compartments represented in the deconvolution analysis and are consistent with transcriptional differences occurring within a broadly comparable cellular context.

Consistent with this interpretation, functional enrichment analyses were performed to characterize the biological programs underlying these composition-independent differences. Upregulated genes in AA relative to CW within G1 included ECM structural components and cell-cycle regulators. Pathway-level enrichment analyses across multiple platforms revealed consistent functional themes centered on two dominant biological programs: extracellular matrix and collagen organization, and mitotic and cytoskeletal organization processes. Downregulated genes in AA relative to CW within G1 showed more limited enrichment, with trends toward ion transport and membrane-associated processes (Supplementary Table S7).

#### 2.4.2 Functional enrichment patterns

Over-representation analysis (Enrichr, GO Biological Process) of genes upregulated in AA within G1 revealed highly significant enrichment for extracellular matrix organization (GO:0030198; adjusted p ≈ 3.1 × 10⁻⁹, odds ratio = 13.8) and a series of mitotic processes including microtubule cytoskeleton organization involved in mitosis (GO:1902850; adjusted p ≈ 8.9 × 10⁻⁸, odds ratio = 26.6), mitotic cytokinesis (GO:0000281; adjusted p ≈ 9.2 × 10⁻⁸), and mitotic spindle midzone assembly (GO:0051256; odds ratio = 95.2) (Supplementary Table S7). topGO analysis confirmed four statistically significant terms: mitotic cytokinesis (adjusted p-value ≈ 0), mitotic spindle midzone assembly (adjusted p-value = 0.0014), cell adhesion (GO:0007155; adjusted p-value = 5.0 × 10⁻⁴), and collagen fibril organization (GO:0030199; adjusted p-value = 5.0 × 10⁻⁴) (Figure 6 and Supplementary Table S7). Network-based analysis using STRING identified two tightly interconnected enriched modules among the AA-upregulated gene set: a mitotic/proliferative module (encompassing chromosome segregation, spindle assembly, and cytokinesis genes) and an extracellular matrix/collagen module (including collagen biosynthesis genes, ECM structural components, and adhesion-related proteins), supported by significant enrichment in both KEGG cell cycle and ECM–receptor interaction pathways (Supplementary Table S7).

**Figure 6.**
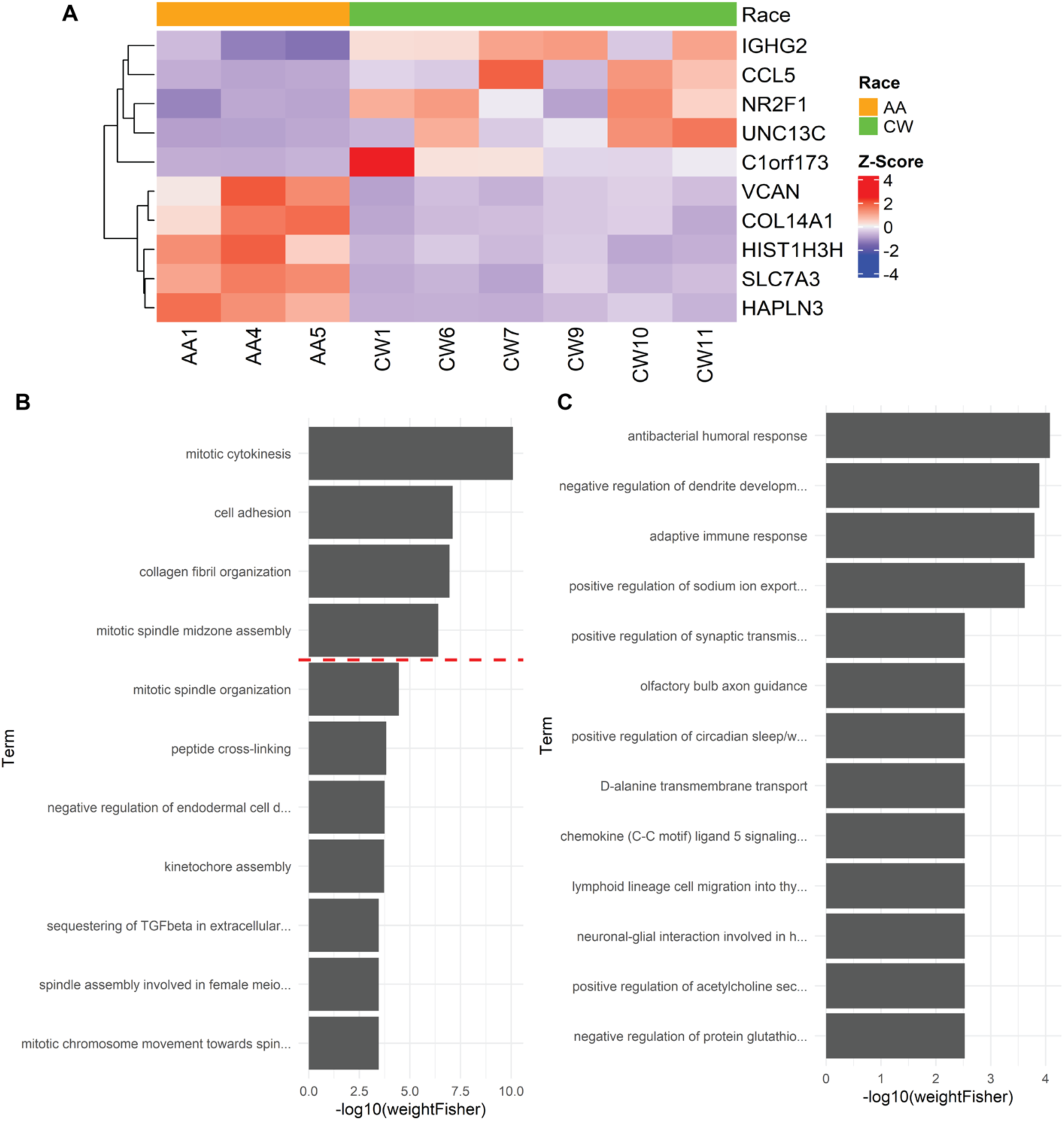
Functional enrichment of transcriptomic differences between population groups within G1. (A) Heatmap of selected differentially expressed genes between AA and CW samples within G1, representing both upregulated (e.g., VCAN, COL14A1, HAPLN3) and downregulated (e.g., IGHG2, CCL5, UNC13C) genes in AA relative to CW. (B) Gene Ontology (GO) Biological Process enrichment analysis (topGO) of genes upregulated in AA relative to CW within G1. Bar plots show the top enriched terms ranked by −log10(weighted Fisher *p*-value), highlighting dominant processes related to mitosis, cytoskeletal organization, and extracellular matrix structure. The dashed red line indicates the significance threshold (weighted Fisher p = 0.05). (C) GO Biological Process enrichment analysis of genes downregulated in AA relative to CW within G1 (i.e., enriched in CW). Enriched terms are ranked by −log10(weighted Fisher *p*-value) and include processes related to immune response, signaling, and cellular communication.

Gene set–based approaches (GSEA, MSigDB C2, C3, C5 collections) showed consistent directional enrichment for ECM organization, collagen-containing pathways, cytoskeletal and spindle-related processes, and cell cycle programs in AA relative to CW within G1, driven by recurrent genes including COL1A1, COL3A1, COL5A1, POSTN, FN1, MKI67, ASPM, CDK1, KIF23, and NUSAP1 (Supplementary Table S7). Although no individual pathway survived multiple testing correction, likely reflecting the limited sample size within this subgroup, the directional patterns were highly consistent with the threshold-based enrichment analyses.

Across all enrichment platforms, extracellular matrix–related and proliferative/cytoskeletal programs were repeatedly identified as the dominant functional categories distinguishing AA from CW samples within G1, providing convergent cross-method support for these biological themes (Table 2 and Supplementary Table S7).

**Table 2.** Summary of functional enrichment results across analytical platforms for genes differentially expressed between population groups within G1. Comparison of enrichment outcomes for major biological themes identified from genes upregulated in African American (AA) relative to Caucasian White (CW) samples within Group 1 (G1). Results are summarized across multiple analytical approaches, including Enrichr, topGO, gene set enrichment analysis (GSEA), and STRING network analysis. “YES (strong)” indicates statistically significant enrichment supported by multiple genes and robust significance thresholds; “YES (moderate)” indicates weaker but consistent enrichment; “directional” indicates consistent trends that did not reach statistical significance after multiple testing correction. The analysis highlights two dominant and consistently supported biological programs, mitotic/cell cycle processes and extracellular matrix (ECM) organization, while ion transport–related processes show weaker and less consistent enrichment across methods

| Method | Mitotic | ECM | Ion Transport |
| --- | --- | --- | --- |
| Enrichr | YES (strong) | YES (strong) | YES (moderate) |
| topGO | YES (strong) | collagen fibril only | NO |
| GSEA | directional only | directional only | directional |
| STRING | YES (strong) | YES (strong) | NO |

#### 2.4.3 Downregulated gene patterns

Genes showing lower expression in AA relative to CW within G1 displayed enrichment for immune-related processes including antibacterial humoral response and adaptive immune response, as well as ion transport terms including positive regulation of sodium ion export (GO:1903276; adjusted p = 0.030, odds ratio = 123.9) (Figure 6C and Supplementary Table S7). STRING returned no statistically significant enrichment for the downregulated gene set, suggesting these genes do not form a tightly interconnected protein interaction network. Given the heterogeneous nature of these terms and the modest statistical support, the downregulated signal should be interpreted as exploratory and hypothesis-generating.

#### 2.4.4 Summary of within-state comparisons

Within G1, population-associated transcriptional differences were substantially more pronounced than in the global cohort analysis, with 191 DEGs identified between AA and CW samples compared to a single gene across the full cohort. These differences were characterized primarily by coordinated upregulation of gene expression programs related to extracellular matrix and proliferative/cytoskeletal programs in AA samples, a pattern consistently supported across multiple enrichment analyses. Downregulated genes showed weaker and more heterogeneous enrichment, with immune-related and ion transport terms that were not consistently supported. No equivalent population-associated enrichment pattern was observed within G2, indicating that these differences are specific to the G1 baseline tissue state.

Notably, UNC13C, the sole gene reaching significance in the full-cohort AA vs CW comparison, was also differentially expressed within G1 (Figure 6A), consistent with the interpretation that the global signal detected in section 2.3.1 is partly driven by the G1 subgroup.

Together, these findings indicate that population-associated molecular differences within this cohort are context-dependent, emerging within a specific baseline transcriptional state characterized by ECM and proliferative programs. While the sample size within this subgroup is limited, the consistency of these patterns across multiple independent enrichment approaches supports the robustness of the observed signal.

### 2.5. Lipidomic Variation across the Full Cohort and Transcriptomic Subgroups

#### 2.5.1 Global lipidomic comparisons between population groups

Untargeted lipidomic profiling was performed using mTIC-normalized intensities from 252 identified lipid species across the full cohort of 30 samples (Figures 7 and 8, and Supplementary Table S8).

**Figure 7.**
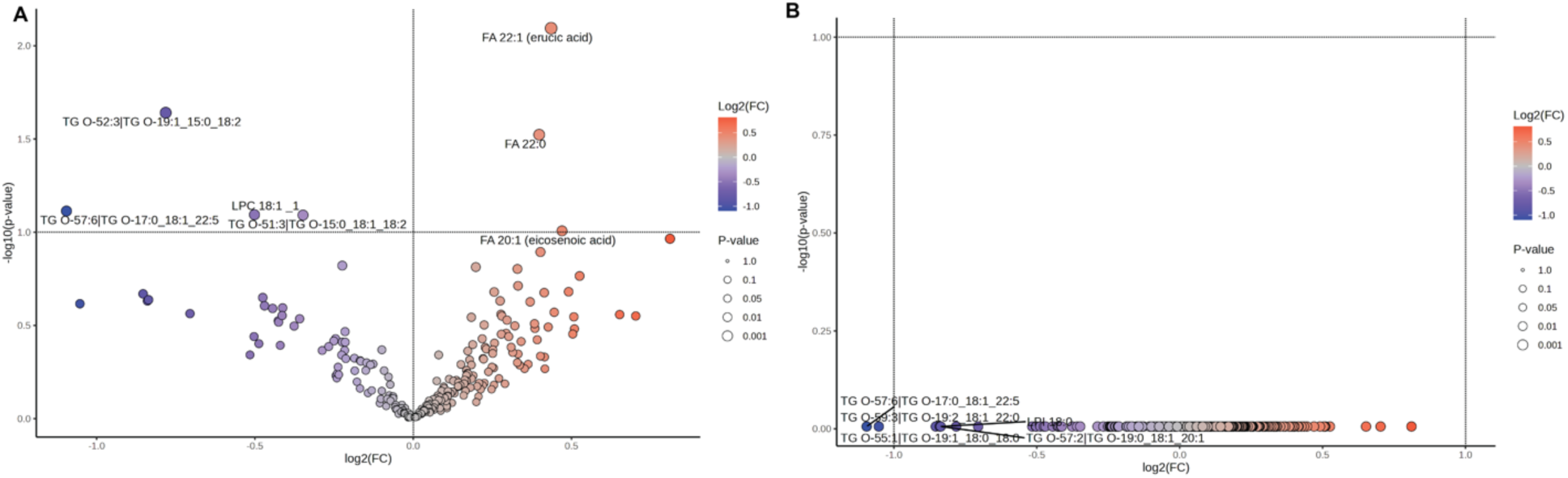
Global lipidomic comparison between population groups (AA vs CW). (A) Volcano plot showing LFC versus −log10(*p*-value) for lipid species comparing AA and CW samples. A small number of lipids show nominal significance (raw *p* ≤ 0.05), but no species remain significant after multiple testing correction. (B) Volcano plot showing LFC versus −log10(adjusted *p*-value). No lipid species meet significance after false discovery rate (FDR) correction.

**Figure 8.**
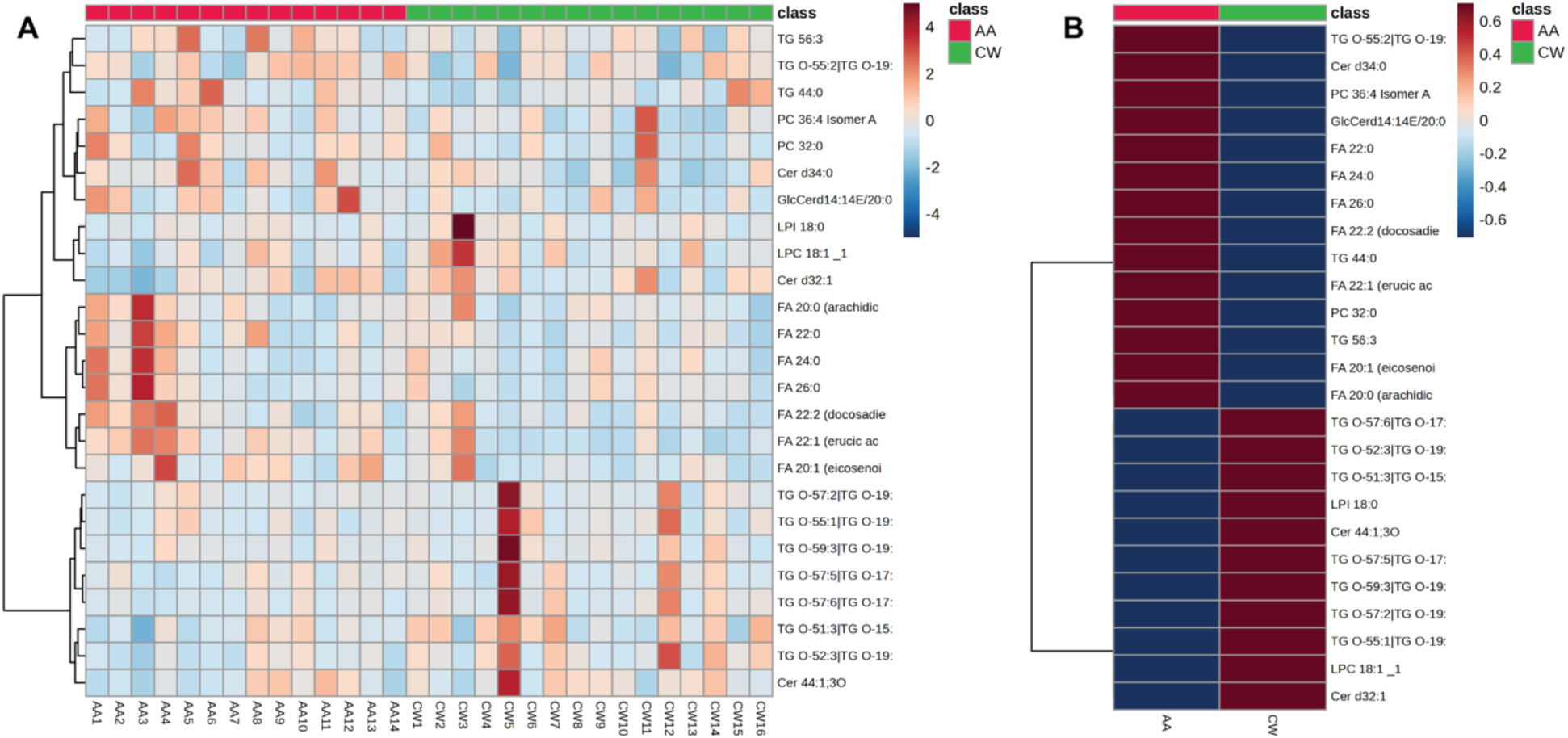
Lipid abundance patterns across population groups (n=30). (A) Heatmap of lipid species showing the largest fold changes between groups across all samples, with rows representing lipids and columns representing individual samples. Values are scaled across samples (Z-score), and samples are annotated by population group (AA and CW). No clear clustering by population group is observed, consistent with the lack of global lipidomic separation. (B) Heatmap summarizing average lipid abundance by population group (AA vs CW) for selected lipid species. Color indicates relative abundance, highlighting modest differences in specific triglycerides and fatty acids between groups without a consistent global pattern.

Univariate comparisons identified three triglycerides, TG 42:3, TG 53:3, and TG 51:2, with nominal differences between groups (raw p ≤ 0.05), all showing higher abundance in CW relative to AA samples. However, no lipid species remained significant after correction for multiple testing. TG O-57:6 and TG O-59:3 showed the largest decreases in AA relative to CW, but these differences were also not statistically significant after FDR correction. Overall, species-level lipidomic variation was dominated by inter-individual variability rather than population group assignment.

Lipids were also grouped into major classes, including triglycerides (TG), fatty acids (FA), ceramides (CER), phospholipids, sphingomyelins (SM), diglycerides (DG), and glycosylated ceramides (GlcCER). Relative lipid class distributions were highly similar between population groups. Triglycerides represented the dominant lipid class in both AA and CW samples, followed by fatty acids, whereas ceramides, phospholipids, sphingomyelins, diglycerides, and glycosylated ceramides contributed smaller proportions. No lipid class differed significantly between groups (Table 3).

**Table 3.** Relative lipid class distribution between population groups across the full cohort (n=30). Comparison of the relative abundance (ratio of total lipid signal) of major lipid classes between African American (AA) and Caucasian White (CW) samples. Ratios represent the fraction of each lipid class normalized to total lipid content per sample. Fold change (AA/CW) reflects relative differences between groups. Statistical significance was assessed using univariate comparisons for each lipid class; no lipid class showed significant differences between groups (*p* > 0.05 for all comparisons). These results indicate that overall lipid class composition is conserved between population groups, with no large-scale redistribution of lipid classes across the cohort

| Lipid Class | AA (ratio) | CW (ratio) | AA/CW Fold Change | p-value |
| --- | --- | --- | --- | --- |
| CER | 0.04575 | 0.04374 | 1.05 | ns |
| DG | 0.00135 | 0.00131 | 1.03 | ns |
| FA | 0.32132 | 0.32490 | 0.99 | ns |
| **GlcCER** | 0.00060 | 0.00046 | 1.30 | ns |
| **Phospholipids** | 0.04974 | 0.04570 | 1.09 | ns |
| **SM** | 0.01047 | 0.00991 | 1.06 | ns |
| **TG** | 0.57077 | 0.57397 | 0.99 | ns |

These findings indicate that the full cohort showed neither robust species-level lipidomic separation nor broad redistribution of major lipid classes by population group.

#### 2.5.2 Lipidomic comparisons between G1 and G2

To determine whether the transcriptomically defined G1 and G2 states were accompanied by broader lipidomic differences, lipid profiles were compared across the 22 samples assigned to these groups. PCA showed substantial overlap between G1 and G2, with PC1 and PC2 explaining 71.2% and 10.7% of total variance, respectively (Figure 9A and Supplementary Figure S3A). Lipids contributing strongly to the PCA structure included FA 18:0, FA 16:0, FA 18:1, and several ether-linked triglycerides, but these features did not independently separate the two groups.

**Figure 9.**
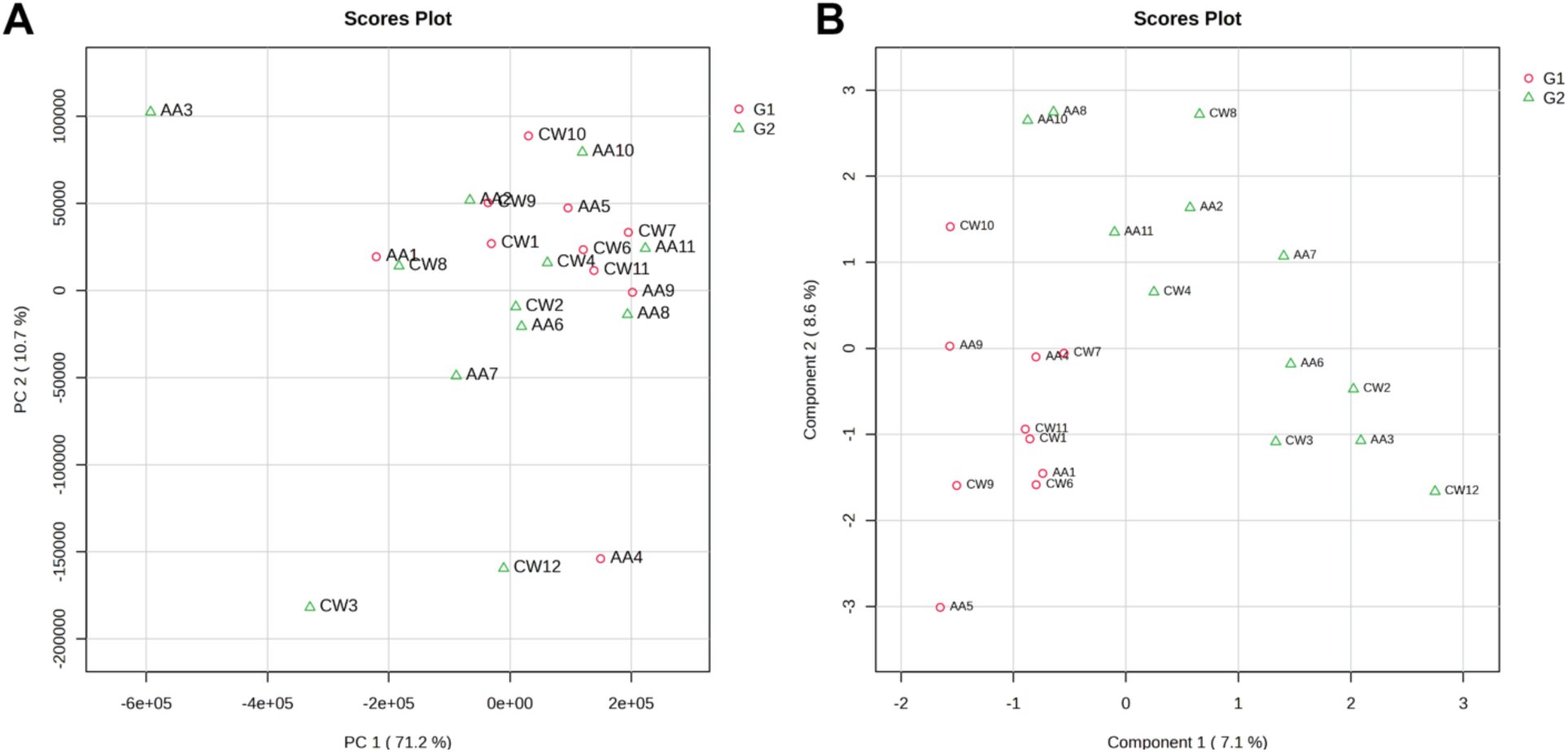
Lipidomic structure relative to transcriptomic group assignment (G1: n=10 and G2: n=12). (A) Principal component analysis (PCA) of lipidomic profiles showing the distribution of samples colored by transcriptomic groups (G1 and G2). PC1 and PC2 explain 71.2% and 10.7% of the variance, respectively. (B) Sparse partial least squares discriminant analysis (sPLS-DA) scores plot of lipidomic profiles colored by transcriptomic group (G1 and G2). Component 1 explains 7.1% of variance.

Supervised sPLS-DA showed partial discrimination between G1 and G2 (Figure 9B). Features contributing to this pattern included ether-linked and polyunsaturated triglycerides, fatty acids, sphingomyelins, ceramides, and selected phospholipids (Supplementary Figure S3B). Because this separation was derived from a supervised analysis and no individual species reached FDR significance, these patterns should be interpreted as exploratory.

Fold-change analysis identified five lipids with FC ≥ 2 in G2 relative to G1: PC 38:4 Isomer C (FC = 3.23), PC 40:5 Isomer A (FC = 2.79), LPE 22:6 (FC = 2.69), PC 36:4 Isomer C (FC = 2.69), and PC 38:5 Isomer A_1 (FC = 2.24), all elevated in G2 (Figure 10E). However, none of these lipids reached FDR significance after multiple testing correction (Figure 10D and F). At the lipid class level, relative distributions remained similar between G1 and G2 with triglycerides dominating both groups and no large-scale class-level redistribution detected (Supplementary Figure S4B and C). Ceramide species appeared nominally among the top candidates in univariate analysis (Cer d34:1, Cer d36:1, Cer d39:1; raw p < 0.05) but none survived FDR correction (Figure 10C, D, and F). These results suggest that lipidomic differences between the two baseline transcriptomic states were subtle relative to overall inter-individual variability (Supplementary Table S9). Relative lipid class distributions were also similar between G1 and G2, with triglycerides dominating both states and no large-scale class-level redistribution detected (Supplementary Figure S4).

**Figure 10.**
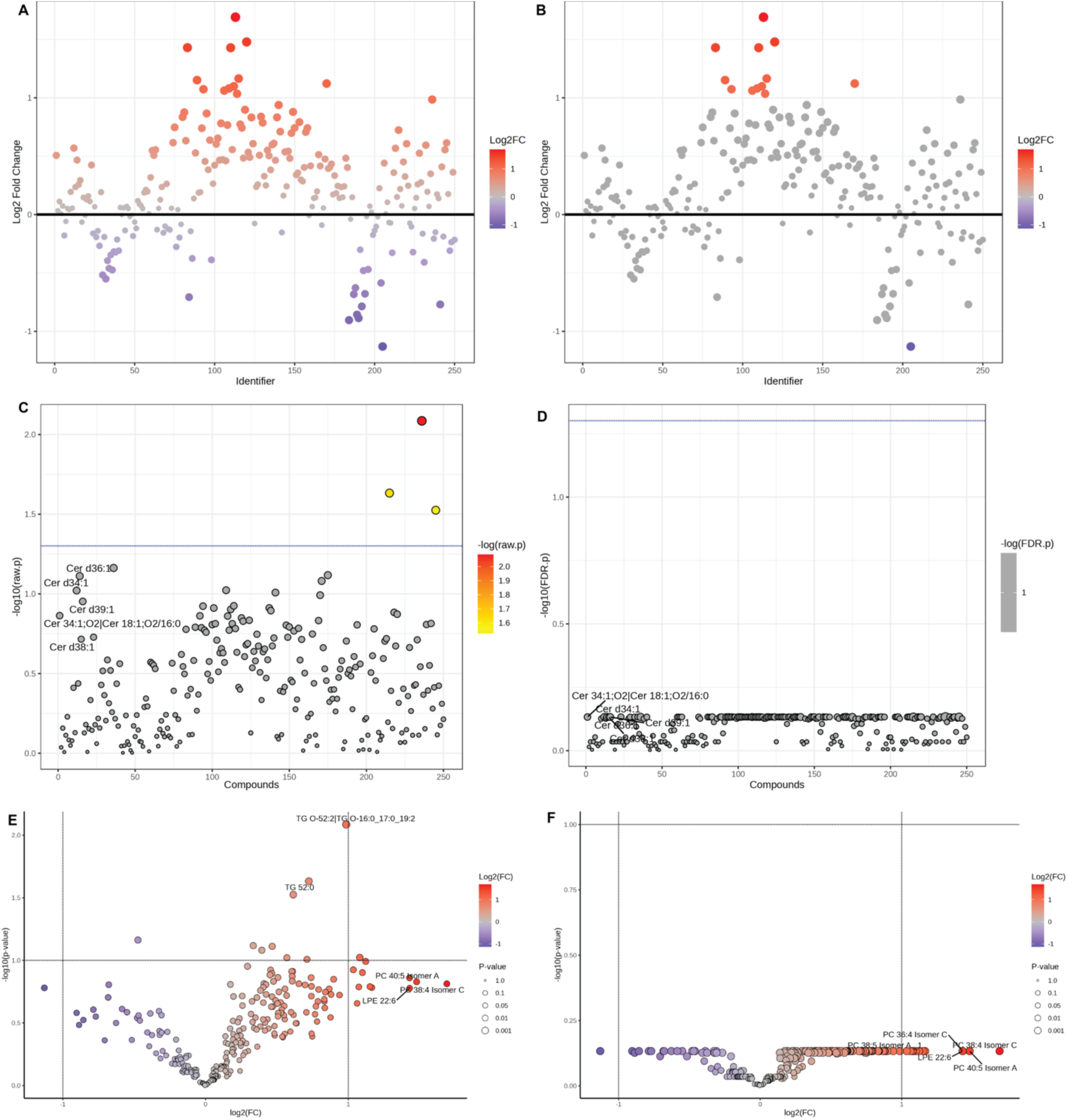
Lipidomic comparison between transcriptomic groups G1 and G2. (A) Scatter plot of LFC for all 252 lipid species comparing G2 relative to G1, ordered by compound identifier and colored by fold change magnitude. Most lipids show modest or no change between groups. (B) Same as (A) with lipids showing |log2FC| ≥ 1 highlighted; remaining lipids are shown in grey. (C) Volcano plot of lipid species based on LFC and −log10(raw *p*-value). Several ceramide species (e.g., Cer d34:1, Cer d36:1, Cer d39:1) and related lipids show nominal significance (raw *p* < 0.05), but overall significance is limited. (D) Volcano plot using FDR-adjusted *p*-values. No lipid species reach statistical significance after multiple testing correction (FDR < 0.05). (E) Volcano plot of LFC versus −log10(raw p-value) for all lipid species, with selected lipids showing the largest fold changes labeled (TG O-52:2, TG 52:0, PC 40:5 Isomer A, PC 38:4 Isomer C, LPE 22:6). (F) Volcano plot based on FDR-adjusted values corresponding to (E), confirming the absence of statistically significant lipid differences after correction.

Together, these results indicate that the two baseline states were strongly resolved at the transcriptomic level but were associated only with subtle and exploratory differences in bulk lipid composition.

#### 2.5.3 Population-associated lipidomic variation within G1

Because population-associated transcriptional differences were detected specifically within G1, lipidomic profiles were also compared between AA and CW samples within this state. In contrast to the transcriptomic analysis, lipidomic profiles did not show robust population-associated separation, and no individual lipid species remained significant after correction for multiple testing. Any species-level or multivariate differences observed within G1 should therefore be considered exploratory.

The absence of significant lipid signals within G1 stands in contrast to the strong transcriptional differences observed in the same samples, indicating that the population-associated variation within G1 is primarily a transcriptional rather than a bulk lipidomic phenomenon.

#### 2.5.4 Summary of lipidomic findings

Across the full cohort, between G1 and G2, and within G1, lipidomic differences were limited relative to the transcriptomic findings. No individual lipid species or major lipid class remained statistically significant after correction for multiple testing, although exploratory multivariate analyses identified modest structure associated with transcriptomic state. These findings indicate that the G1/G2 distinction and the population-associated signal within G1 were primarily transcriptional and were not accompanied by large-scale changes in bulk lipid composition.

### 2.6. Technical and Metadata Evaluation

#### 2.6.1 Assessment of sequencing quality and batch structure

RNA integrity metrics, sequencing parameters, and available facility metadata were reviewed to evaluate potential technical sources of variation. All libraries were prepared in a single batch (June 8–10, 2023) using Illumina TruSeq Stranded Total RNA kits and sequenced on a single NovaSeq 6000 S4 v1.5 flowcell and lane (August 23–24, 2023), eliminating inter-batch variability as an explanation for observed clustering. Per-sample QC metrics from FastQC, STAR, and Picard confirmed consistently high base-call quality (mean Phred ≥ Q30), and no systematic clustering of samples by any available technical variable was observed in principal component space.

#### 2.6.2 Storage category and tissue handling

Storage duration and available handling metadata were examined to determine whether sample age or storage conditions contributed to molecular separation. No consistent association was observed between storage membership in G1 or G2 (Supplementary Figure S1). These observations suggest that long-term storage effects were unlikely to be the primary driver of observed molecular patterns.

#### 2.6.3 Demographic variables and matching strategy

Examination of age and BMI across groups did not reveal segregation with G1 or G2 membership (Figure 1 and Supplementary Figure S1A and S1B). The restricted range of values resulting from matching may have reduced the ability to detect demographic drivers of molecular variation.

#### 2.6.4 Comorbidities and available clinical metadata

Available comorbidity information was reviewed where present. No single comorbidity variable consistently aligned with baseline molecular grouping (data not shown). However, clinical metadata were incomplete for some samples, limiting the ability to fully evaluate all potential clinical contributors to molecular variability. As such, subtle clinical influences cannot be entirely excluded.

## 3. Discussion

African American women experience disproportionately higher breast cancer mortality than Caucasian White women, yet relatively little is known about whether molecular differences are already present in histologically normal breast tissue before disease onset [2,4,5,18–20]. To address this, we performed transcriptomic and lipidomic profiling of histologically normal breast tissue from AA and CW individuals under matched demographic conditions and conceptually integrated the resulting molecular patterns [12]. In this pilot study, the dominant molecular structure of the cohort was defined not by population group assignment, but by two distinct baseline transcriptional states. Population-associated transcriptional differences were limited across the full cohort but became detectable within one of these states, indicating that such variation was context-dependent in the samples examined. These findings suggest that baseline tissue heterogeneity can shape whether population-associated molecular differences are detectable and should therefore be considered a biologically meaningful feature rather than solely an analytical confounder.

The two transcriptional states reflected differences in both tissue composition and transcriptional activity. Cell-type deconvolution indicated that G1 represented a predominantly epithelial-enriched context, whereas G2 was relatively enriched in vascular-associated cells. However, the persistence of substantial transcriptional differences after adjustment for estimated epithelial proportion indicates that cellular composition alone did not account for the separation. Instead, G1 and G2 appear to represent composite tissue states in which differences in cellular representation coexist with distinct transcriptional programs, including increased translational activity in G1 and programs related to endothelial remodeling, cell migration, and metabolism in G2. This interpretation is consistent with previous studies showing that normal breast tissue is molecularly heterogeneous and continuously shaped by variation among epithelial, stromal, vascular, immune, and adipose compartments [1,5,8]. These findings also reinforce the importance of distinguishing intrinsic transcriptional activity from signals produced by differences in tissue architecture when analyzing bulk breast tissue [1,5,8].

Previous work has also identified population-associated variation within histologically normal breast tissue, including differences involving epithelial and vascular-associated programs [1,4,5,7–9]. The present findings extend this literature by suggesting that such differences may not be uniformly distributed across normal tissue. Instead, their detection may depend on the baseline cellular and physiological context in which they occur. Pooling samples representing distinct tissue states may therefore obscure biologically coherent differences that become apparent only after the underlying heterogeneity is recognized. At the same time, because the G1 and G2 states were identified in a modest cohort and have not yet been independently replicated, they should be regarded as provisional biological configurations rather than established categories of normal breast tissue.

The principal population-associated signal was observed within the epithelial-enriched G1 state, where extracellular matrix organization and proliferative/cytoskeletal processes were coordinately increased in AA relative to CW samples. The broad cell-type proportions estimated within G1 were similar between population groups, making it unlikely that this signal was explained solely by differences in the represented epithelial, vascular, stromal, or immune compartments. Nevertheless, the current analysis cannot exclude differences in finer cellular subtypes, cell activation states, spatial organization, or adipocyte abundance. The observed signal is therefore most appropriately interpreted as a transcriptional difference occurring within a broadly comparable cellular context, rather than as definitive evidence of cell-intrinsic or cell-type-specific regulation.

The coexistence of extracellular matrix remodeling and proliferative programs is biologically notable because these processes jointly influence tissue architecture, epithelial behavior, and stromal–epithelial communication [1,8]. The contributing genes span collagen organization, adhesion, cytoskeletal remodeling, spindle assembly, and cell division, consistent with prior work emphasizing the role of the tissue microenvironment in mammary gland biology [1,4,5,7–9]. However, the observed pattern should not be interpreted as evidence of a premalignant state or a complete epithelial-to-mesenchymal transition. Canonical EMT regulators were not strongly enriched, and the findings are better described as a limited, state-associated program of matrix remodeling and proliferation. These processes have been implicated in disease-associated tissue environments, and their coordinated variation in histologically normal tissue is therefore relevant to future studies of susceptibility and disease progression. However, the present cross-sectional data do not establish that they contribute causally to breast cancer risk or outcome. The lack of a comparable detectable signal within G2 further supports a conditional model, although the small subgroup size limits the ability to distinguish true biological absence from insufficient statistical power.

The lipidomic data provided a complementary but less clearly resolved view of the tissue states. Neither population group nor G1/G2 assignment produced robust separation of bulk lipidomic profiles, and no individual lipid species or major lipid class remained significant after correction for multiple testing. Partial separation in supervised analyses and contributions from triglycerides, fatty acids, sphingolipids, ceramides, and phospholipids suggest that subtle lipid variation may accompany the transcriptional states, but these findings remain exploratory. The limited cross-modal concordance may reflect the distinct biological scales captured by the two assays. Transcriptomic profiling can detect coordinated regulatory changes within relatively small cellular compartments, whereas bulk lipidomic measurements in adipose-rich breast tissue may be strongly influenced by overall tissue composition and systemic metabolic state [10,14,21–23]. Variation arising from epithelial or vascular cell populations may therefore be difficult to detect within the dominant lipid signal contributed by adipose tissue. These considerations support the value of integrating transcriptomic and lipidomic data while also recognizing the limitations of direct cross-modal comparison in heterogeneous bulk samples [10,11]. Given the established role of lipid metabolic reprogramming in aggressive breast cancer phenotypes [13–15], cell-type-resolved or spatial lipidomic approaches will be needed to determine whether compartment-specific metabolic differences accompany the transcriptional programs identified here.

Several limitations constrain the interpretation of this study. The cohort was modest, particularly after stratification into G1 and G2, and the matching of age and BMI required selection of a demographically restricted subset that may not represent the broader populations from which the samples were obtained. Available clinical and physiological metadata were incomplete, limiting assessment of factors such as reproductive history, menopausal status, hormonal exposure, medication use, metabolic health, and other environmental or lived influences. Population groups were defined using repository-provided self-identification rather than genomic ancestry, and these categories should not be interpreted as discrete biological classifications. Bulk tissue profiling also cannot determine which cell types produced the observed transcriptional programs or how those cells were spatially organized. Although the deconvolution analysis provided important information about broad epithelial, vascular, stromal, and immune compartments, the reference atlas did not contain an annotated adipocyte population. Differences in adipocyte abundance or adipocyte-associated transcriptional programs therefore could not be evaluated and may contribute to tissue variation not captured by the current analysis. Accordingly, the estimated proportions should be interpreted as relative distributions among the represented epithelial, vascular, stromal, and immune compartments rather than as a complete reconstruction of breast tissue composition. Finally, the absence of an independent validation cohort and matched functional or tumor data limits assessment of the reproducibility and disease relevance of the identified states.

Future studies should first determine whether the G1 and G2 configurations recur in larger and independently collected normal breast tissue cohorts. Expanded studies should integrate genomic ancestry with detailed clinical, hormonal, reproductive, metabolic, socioeconomic, and environmental information to evaluate the sources of the observed variation [1–3,5,8,24]. Single-cell and spatial transcriptomic approaches will be particularly important for identifying the epithelial, stromal, vascular, or immune populations responsible for the ECM and proliferative signals and for determining whether the relevant differences reflect cell-intrinsic regulation, altered cellular states, or changes in intercellular organization. Parallel cell-type-resolved or spatial lipidomic analyses may reveal metabolic differences that are obscured in bulk adipose-rich tissue [1–3,5,8,24]. Studies incorporating matched normal and tumor tissue, or longitudinal clinical follow-up, will ultimately be required to determine whether these baseline programs are associated with tumor subtype, disease characteristics, or clinical trajectory, or whether they remain within the range of normal physiological variation.

In summary, this study suggests that molecular variation in histologically normal breast tissue is structured and conditional rather than uniformly distributed across population groups. The dominant axis of variation reflected baseline tissue state, while a coordinated extracellular matrix and proliferative/cytoskeletal program emerged specifically within the epithelial-enriched G1 context. These findings do not establish a direct link to disease susceptibility, but they provide a framework for investigating how baseline cellular composition, tissue architecture, and regulatory activity may be shaped by genetic ancestry and environmental or physiological history. Larger, deeply characterized, and cell-type-resolved studies will be required to determine whether these context-dependent differences have functional consequences for breast cancer susceptibility, tumor characteristics, progression, or population-associated disparities [1–9,25].

## 4. Methods

### 4.1 Study Cohort and Sample Metadata

#### 4.1.1 Tissue source and cohort structure

Histologically normal human breast tissue samples were obtained from a breast tissue repository associated with Susan G. Komen Tissue Bank at the Indiana University Simon Cancer Center (through a material transfer agreement #186667). The study cohort consisted of 30 samples derived from individuals classified by the repository metadata as African American (AA; n = 14) or Caucasian White (CW; n = 16).

Throughout the manuscript, the term “self-identified racial groups” refers to the donor classifications provided by the repository, while “population groups” is used in the analytical description of aggregated comparisons. To reduce major demographic confounding effects, samples were selected to achieve approximate matching of age and body mass index (BMI) across groups. Due to availability constraints in the repository, matching required selecting AA samples within a narrower BMI range to ensure overlap with CW samples. As a result, the analyzed dataset represents a demographically constrained subset rather than a representative population sample.

Available metadata included donor age, BMI, sample identifiers, tissue cellularity annotations, and sample storage metadata. These variables were evaluated during downstream analyses to assess potential non-biological drivers of transcriptomic variation.

#### 4.1.2 Metadata processing and cohort annotation

Sample metadata were curated and organized into a unified dataset linking RNA-seq identifiers, lipidomics identifiers, and repository metadata. Variables examined during downstream evaluation included age, BMI, storage duration, tissue cellularity annotations, and reported comorbidities. To evaluate potential demographic or technical biases, these metadata variables were later overlaid on transcriptomic principal component analysis (PCA) plots and used in exploratory statistical comparisons between groups.

### 4.2 RNA Sequencing

#### 4.2.1 Library preparation

RNA sequencing libraries were prepared at the University of California, Irvine Genomics High-Throughput Facility (GHTF). All 30 samples were processed together in a single library preparation batch between June 8 and June 10, 2023, using the Illumina TruSeq Stranded Total RNA library preparation kit. The initial experimental design involved poly-A mRNA enrichment; however, because the submitted RNA samples exhibited substantial degradation, the library preparation strategy was modified to use ribosomal RNA depletion following consultation between the sequencing facility and investigators. This approach allowed retention of fragmented transcripts while minimizing rRNA contamination. All libraries were prepared by the same technician, thereby reducing variability due to operator effects.

#### 4.2.2 Sequencing

Sequencing was performed on an Illumina NovaSeq 6000 platform using an S4 v1.5 flow cell. All libraries were sequenced together in a single sequencing run (run identifier HFWT3DSX7) on August 23–24, 2023, and multiplexed within lane 1. Demultiplexing was carried out by the sequencing facility using unique dual indices assigned to each sample. Sequencing yielded between approximately 16.3 and 31.9 million paired reads per sample. Run-level sequencing metrics indicated high overall sequencing quality, with >93% bases exceeding Q30 across the run. Because all libraries were prepared and sequenced together, the dataset was generated under a single-batch design, minimizing typical cross-batch sequencing artifacts.

#### 4.2.3 RNA-seq data processing and quality control

Raw FASTQ files were processed using the nf-core/rnaseq pipeline (v3.14.0) [26]. Standard pipeline modules were used for quality assessment, alignment, transcript quantification, and generation of gene-level count matrices. Quality metrics from the pipeline were aggregated using MultiQC, [27] enabling evaluation of sequencing quality, mapping statistics, duplication rates, and library complexity.

Samples exhibiting poor sequencing quality were excluded prior to downstream analysis. Exclusion criteria included high duplication rates (>50%) and low proportions of properly paired reads (<30%), consistent with reduced library complexity and unreliable transcript quantification (Supplementary Table S1, second sheet). A total of 8 samples were excluded on this basis (Table 1; Group = NA: CW5, AA12, CW13, CW14, CW15, CW16, AA13, AA14). After quality control filtering, 22 samples remained for downstream transcriptomic analysis, comprising 10 African American (AA) and 12 Caucasian White (CW) samples. These samples were distributed across Group 1 (G1; n = 10) and Group 2 (G2; n = 12), with both population groups represented in each cluster. Excluded samples were distributed across population groups and did not show systematic bias toward a specific group.

Because all libraries were processed in a single sequencing run, sequencing batch effects were not modeled explicitly in downstream statistical analyses. Instead, potential technical effects were evaluated through metadata overlays and inspection of facility-provided quality metrics.

### 4.3 Transcriptomic Analysis

#### 4.3.1 Principal component analysis and identification of transcriptomic states

Normalized gene expression matrices derived from the RNA-seq workflow were used for unsupervised dimensionality reduction [28,29]. Principal component analysis (PCA), using the R package pcaExplorer (v3.0.0) [30], was applied to identify major axes of transcriptional variation across samples. Initial PCA revealed the presence of two major transcriptional clusters that did not correspond to the repository-defined racial group assignments; these clusters were designated Group 1 (G1) and Group 2 (G2) and served as a framework for subsequent analyses. To confirm this structure independently, t-distributed stochastic neighbor embedding (t-SNE) [31] was performed using the same expression matrix. In later stages of the analysis, samples that did not cluster clearly with either transcriptional group were excluded from the refined comparison to sharpen separation between the dominant transcriptomic states; the refined PCA used for final analyses therefore reflects a filtered subset of the dataset.

#### 4.3.2 Differential gene expression analysis

Differential gene expression analysis was performed using the DESeq2 (v1.46.0)[32] package on raw gene-level counts. To facilitate direct pairwise comparisons between populations or groups, three comparisons were performed: (i) AA vs CW across the full cohort, (ii) G1 vs G2 transcriptomic states across both population groups, and (iii) AA vs CW within the G1 subgroup. For the global AA vs CW comparison, significance thresholds were defined as an adjusted p-value ≤ 0.1 and |LFC| ≥ 1; this relaxed FDR threshold was applied because of the limited signal detected in the global comparison. For the G1 vs G2 and AA vs CW within G1 comparisons, more stringent thresholds were applied: adjusted p-value ≤ 0.05 an |LFC| ≥ 1. Multiple testing correction was performed using the Benjamini–Hochberg procedure. Results were visualized using volcano plots, MA plots, and expression heatmaps. To assess the contribution of cellular composition to the G1/G2 transcriptional separation, a composition-corrected differential expression model was additionally applied; details of this analysis are described in Section 4.7.4.

### 4.4 Functional Enrichment and Network Analysis

#### 4.4.1 Enrichr over-representation analysis

Gene Ontology Biological Process enrichment analysis was performed using Enrichr [33] on sets of significantly upregulated and downregulated genes. Enrichment significance was determined using adjusted p-values derived from Fisher’s exact test. Terms with adjusted p-values < 0.05 were considered significant.

#### 4.4.2 Gene set enrichment analysis

Gene set enrichment analysis (GSEA) [34] was performed using the fgsea (v1.32.2) [35] implementation on ranked gene lists derived from differential expression statistics. Analyses were conducted using selected gene sets from the Molecular Signatures Database (MSigDB) [36], including collections C2, C3, and C5. Pathway enrichment was interpreted using normalized enrichment scores (NES) and adjusted p-values.

#### 4.4.3 topGO analysis

Gene Ontology enrichment was also evaluated using the topGO (v2.58.0) [37] package with the weight01 algorithm and Fisher’s exact test. This approach accounts for the hierarchical structure of GO terms and provides a conservative complement to standard over-representation analyses.

#### 4.4.4 Protein interaction network analysis

Protein interaction networks were examined using STRING (version 12.0) [38] with a minimum interaction confidence threshold of 0.7. Network enrichment analysis was used to identify functional interaction clusters among differentially expressed genes, particularly those related to extracellular matrix organization, cell cycle regulation, and adhesion pathways.

### 4.5 Lipidomic Profiling and Analysis

#### 4.5.1 Lipidomic data generation

Lipid extraction and analysis were conducted using a biphasic lipid extraction method at the UC Davis West Coast Metabolomics Center. Lipids were separated using a C18-based hybrid bridged column with a ternary water/acetonitrile/isopropanol gradient and analyzed on a ThermoFisher Scientific Q-Exactive HF mass spectrometer (Thermo Fisher Scientific, Bremen, Germany) with heated electrospray ionization (HESI-MS). For ionization, the HESI-MS operated at a spray voltage of 2.5 kV (positive mode) and 3.5 kV (negative mode). The capillary temperature was set to 290 °C, with a sheath gas flow rate of 60 arbitrary units, auxiliary gas set to 18 arbitrary units at a probe temperature of 475 °C, and a curtain gas setting of 3.5 arbitrary units.

For data-dependent acquisition (DDA), precursor ions were selected using a 4 Da isolation width and fragmented with a collision energy of 25 eV for both positive and negative ion modes. MS/MS spectra were acquired in data-dependent mode at a resolution of 10,000 (ESI+) and 20,000 (ESI−) using an Agilent 6530 and 6550 QTOF MS (Agilent Technologies, Santa Clara, CA, USA), respectively. The spectral acquisition speed was set to 2 spectra per second, and precursor selection was dynamically excluded after two occurrences to minimize redundant fragmentation.

Lipid annotations were processed using MS-DIAL v4.90, with precursor mass errors < 10 mDa, retention time matching, and MS/MS matching to corresponding lipid classes [39]. Lipid annotations were assigned based on total carbon number and degree of unsaturation, with structural assignments following the convention X:Y, where X denotes the total number of carbon atoms and Y represents the number of double bonds (e.g., TAG 51:2). Lipid intensities were sum-normalized (mTIC) by representing each lipid as a fraction of total lipids within the sample, scaled by the median intensity for each treatment group [40–42]. All reported lipid identifications are consistent with Supplementary Table S6, which provides detailed annotations from our lipidomics analysis.

#### 4.5.2 Lipidomic data preprocessing

Analyses were performed on the table of identified lipid species, comprising 252 annotated lipid features. Lipid abundance values were analyzed using mTIC-normalized intensities, consistent with the analytical framework used in our recent lipidomics studies [29,43,44]. Lipids were grouped into seven major classes: triglycerides (TG), fatty acids (FA), ceramides (CER), phospholipids, sphingomyelins (SM), diglycerides (DG), and glycosylated ceramides (GlcCER). Class-level summaries were calculated by aggregating normalized intensities across lipid species within each class.

#### 4.5.3 Lipidomic statistical analysis

Lipidomic comparisons were conducted for three contrasts: AA vs CW across the full cohort, G1 vs G2 transcriptomic groups, and AA vs CW within the G1 subgroup using MetaboAnalyst (version 6.0) [45]. Univariate analyses included fold-change calculations, two-sided unpaired t-tests, and volcano plot visualization. P-values were corrected using the Benjamini–Hochberg procedure (FDR <= 0.05). Because few lipid species remained significant after FDR correction, lipidomic results were interpreted primarily as exploratory signals rather than definitive drivers of group separation.

Lipid abundance data were further visualized using boxplots and heatmaps to represent total and proportional lipid class abundances across samples and conditions. Heatmaps were generated using hierarchical clustering with Euclidean distance and Ward’s linkage method.

Multivariate analyses were performed using principal component analysis (PCA) to identify major sources of variance, with results visualized using scree plots and two-dimensional score plots, and sparse partial least squares discriminant analysis (sPLS-DA) as a supervised method to identify lipid features contributing to group discrimination. Together, these analyses were used to evaluate whether lipidomic variation corresponded to transcription-defined sample states and to assess the extent to which lipid profiles reflected underlying molecular structure.

### 4.6 Metadata Evaluation

To determine whether the observed transcriptomic separation (G1 vs G2) could be explained by demographic or technical variables, metadata factors were examined using both statistical testing and PCA overlays. Variables evaluated included age, BMI, tissue storage duration, tissue cellularity annotations, and selected comorbidity variables. Continuous variables were compared using appropriate parametric or nonparametric tests, while categorical variables were evaluated using Fisher’s exact test. Because RNA libraries were prepared and sequenced in a single batch, sequencing batch was not included as a covariate in statistical models.

### 4.7 Cell-type Deconvolution

#### 4.7.1 Reference dataset and deconvolution framework

Cell-type composition of bulk RNA-seq samples was estimated using MuSiC (Multi-Subject Single-Cell deconvolution), a reference-based deconvolution framework that uses a single-cell RNA-seq reference to estimate cell-type proportions in bulk transcriptomic data [17]. Briefly, MuSiC fits a weighted non-negative least squares model that accounts for cross-subject variability in the reference dataset to produce robust proportion estimates across samples.

Deconvolution was applied to the normalized bulk RNA-seq expression matrix of retained samples (n = 22) using default MuSiC parameters. The reference single-cell RNA-seq dataset (breast_atlas.h5ad; https://datasets.cellxgene.cziscience.com/2df9adc8-fefc-4a96-9d80-0a7c6b4576e1.h5ad; Human Breast Cell Atlas - global; [46]) was derived from normal human mammary gland tissue and comprised the following annotated cell types: luminal epithelial cells, basal-myoepithelial cells, luminal adaptive secretory precursor cells, luminal hormone-sensing cells, blood vessel endothelial cells, lymphatic endothelial cells, fibroblasts, mural cells, perivascular cells, and leukocytes [17]. The reference dataset did not contain an annotated adipocyte population. Although the reference-construction pipeline included a category for identifying and collapsing adipocyte annotations, no adipocyte population was present in the source atlas or the final reference object used for deconvolution. Consequently, the estimates reflect the relative contributions of epithelial, vascular, stromal, and immune compartments and do not provide information regarding adipocyte abundance.

#### 4.7.2 Cell-type category collapsing

Individual cell-type estimates were aggregated into four broad categories to reduce multiple testing burden and improve statistical power: epithelial (luminal epithelial, basal-myoepithelial, and secretory precursor cells), vascular (blood vessel and lymphatic endothelial cells), stromal (fibroblasts, with mural and perivascular cells retained in the full output but contributing negligibly to collapsed estimates), and immune (leukocytes). Four cell types, luminal hormone-sensing cells, generic mammary gland epithelial cells, mural cells, and perivascular cells, showed zero or near-zero estimated proportions across all samples and contributed negligibly to the collapsed categories. Full individual cell-type results are provided in Supplementary Table S3.

#### 4.7.3 Statistical comparisons of cell-type proportions

Differences in collapsed cell-type proportions were assessed using two-sided Wilcoxon rank-sum tests for two comparisons: G1 versus G2 (n = 10 vs n = 12) and AA versus CW within G1 (n = 4 vs n = 6). P-values were corrected for multiple testing within each comparison using the Benjamini-Hochberg false discovery rate procedure applied across the four collapsed categories. Statistical analyses were performed in R using the base stats package.

#### 4.7.4 Composition-corrected differential expression analysis

To evaluate whether the G1/G2 transcriptional separation persisted after accounting for cellular composition, a composition-corrected differential expression model was applied. Estimated epithelial cell-type proportions were scaled (mean-centered and divided by standard deviation) and included as a continuous covariate in the DESeq2 design formula alongside the primary grouping variable (design: ∼ epithelial_scaled + group). The same significance thresholds were applied as in the primary analysis (adjusted p-value ≤ 0.05 and |LFC| ≥ 1). GO Biological Process enrichment analysis of composition-independent DEGs was performed using both Enrichr and STRING as described in sections 4.4.1 and 4.4.4 respectively. Results are provided in Supplementary Table S4.

### 4.8 Statistical Analysis and Visualization

Statistical analyses were performed using R-based workflows consistent with those used in our recent transcriptomic and lipidomic studies including R packages such as tidyverse (v2.0.0), biomaRt (v2.62.1), Rtsne (v0.17), BinfTools (v1.0.0), and Rgraphviz (v2.50.0). For omics-scale analyses, p-values were adjusted for multiple testing using the Benjamini–Hochberg procedure. Visualization of transcriptomic and lipidomic data included PCA plots, volcano plots, heatmaps, and enrichment dot plots generated using standard R plotting libraries.

## Supplementary Materials

Supplementary Tables S1-S9 and Supplementary Figures S1-S4.

## Author Contributions

Conceptualization, NN and OVR; Human tissue sample acquisition, OVR, YZ, NN; Data curation, DC, and NN; Formal analysis, WDH, KS, YC, DC, GH, and NN; Funding acquisition, NN, OVR; Methodology, WDH, KS, DC, OVR and NN; Resources, OVR, NN; Writing – original draft, WDH, KS, DC, GH, OVR, and NN; Writing – review & editing, WDH, KS, DC, GH, YZ, OVR, and NN.

## Data Availability Statement

All data reported are provided in the text and supplemental materials. All data reported are provided in the text and supplemental materials. The raw sequencing data are hosted at NCBI (GSE333857). The code used is provided at https://github.com/WDHulsy/CSUF-nikolaidis-breast-tissue/tree/main.

## Acknowledgments

The authors thank the Susan G. Komen Tissue Bank at the Indiana University Simon Cancer Center for providing the breast tissue samples used in this study. Lipidomic profiling was performed at the UC Davis West Coast Metabolomics Center directed by Dr. Oliver Fiehn, whose expertise and resources were essential to this work.

## References Cited

1. Polyak, K.; Kalluri, R. The role of the microenvironment in mammary gland development and cancer. Cold Spring Harb Perspect Biol 2010, 2, a003244, doi:10.1101/cshperspect.a003244.

2. Kim, G.; Pastoriza, J.M.; Condeelis, J.S.; Sparano, J.A.; Filippou, P.S.; Karagiannis, G.S.; Oktay, M.H. The Contribution of Race to Breast Tumor Microenvironment Composition and Disease Progression. Front Oncol 2020, 10, 1022, doi:10.3389/fonc.2020.01022.

3. Andrade de Oliveira, K.; Sengupta, S.; Yadav, A.K.; Clarke, R. The complex nature of heterogeneity and its roles in breast cancer biology and therapeutic responsiveness. Front Endocrinol (Lausanne) 2023, 14, 1083048, doi:10.3389/fendo.2023.1083048.

4. Bhat-Nakshatri, P.; Gao, H.; Sheng, L.; McGuire, P.C.; Xuei, X.; Wan, J.; Liu, Y.; Althouse, S.K.; Colter, A.; Sandusky, G.;, et al. A single-cell atlas of the healthy breast tissues reveals clinically relevant clusters of breast epithelial cells. Cell Rep Med 2021, 2, 100219, doi:10.1016/j.xcrm.2021.100219.

5. Nakshatri, H.; Anjanappa, M.; Bhat-Nakshatri, P. Ethnicity-Dependent and -Independent Heterogeneity in Healthy Normal Breast Hierarchy Impacts Tumor Characterization. Scientific Reports 2015, 5, 13526, doi:10.1038/srep13526.

6. Danforth, D.N., Jr. Genomic Changes in Normal Breast Tissue in Women at Normal Risk or at High Risk for Breast Cancer. Breast Cancer (Auckl) 2016, 10, 109–146, doi:10.4137/bcbcr.S39384.

7. Parsons, A.; Sauras Colón, E.; Manjunath, M.; Zhang, H.; Chen, J.; Spasic, M.; Koca, B.; Binboga Kurt, B.; Freedman, R.A.; Mittendorf, E.A.;, et al. Cell populations in human breast cancers are molecularly and biologically distinct with age. Nature Aging 2025, 5, 2546–2563, doi:10.1038/s43587-025-00984-1.

8. Johansson, H.J.; Socciarelli, F.; Vacanti, N.M.; Haugen, M.H.; Zhu, Y.; Siavelis, I.; Fernandez-Woodbridge, A.; Aure, M.R.; Sennblad, B.; Vesterlund, M.;, et al. Breast cancer quantitative proteome and proteogenomic landscape. Nature Communications 2019, 10, 1600, doi:10.1038/s41467-019-09018-y.

9. Asselin-Labat, M.L.; Vaillant, F.; Sheridan, J.M.; Pal, B.; Wu, D.; Simpson, E.R.; Yasuda, H.; Smyth, G.K.; Martin, T.J.; Lindeman, G.J.;, et al. Control of mammary stem cell function by steroid hormone signalling. Nature 2010, 465, 798–802, doi:10.1038/nature09027.

10. Taware, R.; More, T.H.; Bagadi, M.; Taunk, K.; Mane, A.; Rapole, S. Lipidomics investigations into the tissue phospholipidomic landscape of invasive ductal carcinoma of the breast. RSC Adv 2020, 11, 397–407, doi:10.1039/d0ra07368g.

11. Song, H.; Tang, X.; Liu, M.; Wang, G.; Yuan, Y.; Pang, R.; Wang, C.; Zhou, J.; Yang, Y.; Zhang, M.;, et al. Multi-omic analysis identifies metabolic biomarkers for the early detection of breast cancer and therapeutic response prediction. iScience 2024, 27, 110682, doi:10.1016/j.isci.2024.110682.

12. Cavill, R.; Jennen, D.; Kleinjans, J.; Briede, J.J. Transcriptomic and metabolomic data integration. Brief Bioinform 2016, 17, 891–901, doi:10.1093/bib/bbv090.

13. Perrotti, F.; Rosa, C.; Cicalini, I.; Sacchetta, P.; Del Boccio, P.; Genovesi, D.; Pieragostino, D. Advances in Lipidomics for Cancer Biomarkers Discovery. Int J Mol Sci 2016, 17, doi:10.3390/ijms17121992.

14. Wright, H.J.; Hou, J.; Xu, B.; Cortez, M.; Potma, E.O.; Tromberg, B.J.; Razorenova, O.V. CDCP1 drives triple-negative breast cancer metastasis through reduction of lipid-droplet abundance and stimulation of fatty acid oxidation. Proc Natl Acad Sci U S A 2017, 114, E6556–e6565, doi:10.1073/pnas.1703791114.

15. Park, J.H.; Vithayathil, S.; Kumar, S.; Sung, P.L.; Dobrolecki, L.E.; Putluri, V.; Bhat, V.B.; Bhowmik, S.K.; Gupta, V.; Arora, K.;, et al. Fatty Acid Oxidation-Driven Src Links Mitochondrial Energy Reprogramming and Oncogenic Properties in Triple-Negative Breast Cancer. Cell Rep 2016, 14, 2154–2165, doi:10.1016/j.celrep.2016.02.004.

16. Camarda, R.; Zhou, A.Y.; Kohnz, R.A.; Balakrishnan, S.; Mahieu, C.; Anderton, B.; Eyob, H.; Kajimura, S.; Tward, A.; Krings, G.;, et al. Inhibition of fatty acid oxidation as a therapy for MYC-overexpressing triple-negative breast cancer. Nat Med 2016, 22, 427–432, doi:10.1038/nm.4055.

17. Wang, X.; Park, J.; Susztak, K.; Zhang, N.R.; Li, M. Bulk tissue cell type deconvolution with multi-subject single-cell expression reference. Nature Communications 2019, 10, 380, doi:10.1038/s41467-018-08023-x.

18. Ribeiro, H.F.; Pelloso, F.C.; Fonseca, B.S.D.; Camparoto, C.W.; Carvalho, M.D.B.; Marques, V.D.; Bitencourt, M.R.; Stevanato, K.P.; Borba, P.B.; Borghesan, D.H.P.;, et al. Racial and Socioeconomic Disparity in Breast Cancer Mortality: A Systematic Review and Meta-Analysis. Cancers (Basel) 2025, 17, doi:10.3390/cancers17101641.

19. Tao, L.; Gomez, S.L.; Keegan, T.H.; Kurian, A.W.; Clarke, C.A. Breast Cancer Mortality in African-American and Non-Hispanic White Women by Molecular Subtype and Stage at Diagnosis: A Population-Based Study. Cancer Epidemiol Biomarkers Prev 2015, 24, 1039–1045, doi:10.1158/1055-9965.Epi-15-0243.

20. Wannaphut, C.; Ponvilawan, B.; Iwase, T.; Takahashi, Y.; Wang, X.; Ueno, N.T. Racial and ethnic disparities in survival outcomes among patients with inflammatory breast cancer: a systematic review and meta-analysis. npj Breast Cancer 2025, 12, 14, doi:10.1038/s41523-025-00879-9.

21. Zhang, F.; Du, G. Dysregulated lipid metabolism in cancer. World J Biol Chem 2012, 3, 167–174, doi:10.4331/wjbc.v3.i8.167.

22. Vidavsky, N.; Kunitake, J.; Diaz-Rubio, M.E.; Chiou, A.E.; Loh, H.C.; Zhang, S.; Masic, A.; Fischbach, C.; Estroff, L.A. Mapping and Profiling Lipid Distribution in a 3D Model of Breast Cancer Progression. ACS Cent Sci 2019, 5, 768–780, doi:10.1021/acscentsci.8b00932.

23. Purwaha, P.; Gu, F.; Piyarathna, D.W.B.; Rajendiran, T.; Ravindran, A.; Omilian, A.R.; Jiralerspong, S.; Das, G.; Morrison, C.; Ambrosone, C.;, et al. Unbiased Lipidomic Profiling of Triple-Negative Breast Cancer Tissues Reveals the Association of Sphingomyelin Levels with Patient Disease-Free Survival. Metabolites 2018, 8, doi:10.3390/metabo8030041.

24. Calvo, I.; Maimó-Barceló, A.; Garate, J.; Bestard-Escalas, J.; Scrimini, S.; Sauleda, J.; Cosío, B.G.; Fernández, J.A.; Barceló-Coblijn, G. Challenges and Advantages of Using Spatially Resolved Lipidomics to Assess the Pathological State of Human Lung Tissue. Cancers (Basel) 2025, 17, doi:10.3390/cancers17132160.

25. Gadaleta, E.; Fourgoux, P.; Pirró, S.; Thorn, G.J.; Nelan, R.; Ironside, A.; Rajeeve, V.; Cutillas, P.R.; Lobley, A.E.; Wang, J.;, et al. Characterization of four subtypes in morphologically normal tissue excised proximal and distal to breast cancer. npj Breast Cancer 2020, 6, 38, doi:10.1038/s41523-020-00182-9.

26. Ewels, P.A.; Peltzer, A.; Fillinger, S.; Patel, H.; Alneberg, J.; Wilm, A.; Garcia, M.U.; Di Tommaso, P.; Nahnsen, S. The nf-core framework for community-curated bioinformatics pipelines. Nature Biotechnology 2020, 38, 276–278, doi:10.1038/s41587-020-0439-x.

27. Ewels, P.; Magnusson, M.; Lundin, S.; Käller, M. MultiQC: summarize analysis results for multiple tools and samples in a single report. Bioinformatics 2016, 32, 3047–3048, doi:10.1093/bioinformatics/btw354.

28. Reinschmidt, A.; Solano, L.; Chavez, Y.; Hulsy, W.D.; Nikolaidis, N. Transcriptomics Unveil Canonical and Non-Canonical Heat Shock-Induced Pathways in Human Cell Lines. Int J Mol Sci 2025, 26, doi:10.3390/ijms26031057.

29. Solano, L.E.; Keshet, U.; Reinschmidt, A.; Chavez, Y.; Hulsy, W.D.; Fiehn, O.; Nikolaidis, N. Dynamic Lipidome Reorganization in Response to Heat Shock Stress. Int J Mol Sci 2025, 26, doi:10.3390/ijms26072843.

30. Marini, F.; Binder, H. pcaExplorer: an R/Bioconductor package for interacting with RNA-seq principal components. BMC Bioinformatics 2019, 20, 331, doi:10.1186/s12859-019-2879-1.

31. van der Maaten, L.; Hinton, G. Visualizing Data using t-SNE. Journal of Machine Learning Research 2008, 9, 2579--2605.

32. Love, M.I.; Huber, W.; Anders, S. Moderated estimation of fold change and dispersion for RNA-seq data with DESeq2. Genome Biol 2014, 15, 550, doi:10.1186/s13059-014-0550-8.

33. Xie, Z.; Bailey, A.; Kuleshov, M.V.; Clarke, D.J.B.; Evangelista, J.E.; Jenkins, S.L.; Lachmann, A.; Wojciechowicz, M.L.; Kropiwnicki, E.; Jagodnik, K.M.;, et al. Gene Set Knowledge Discovery with Enrichr. Current Protocols 2021, 1, e90, 10.1002/cpz1.90.

34. Subramanian, A.; Tamayo, P.; Mootha, V.K.; Mukherjee, S.; Ebert, B.L.; Gillette, M.A.; Paulovich, A.; Pomeroy, S.L.; Golub, T.R.; Lander, E.S.;, et al. Gene set enrichment analysis: a knowledge-based approach for interpreting genome-wide expression profiles. Proc Natl Acad Sci U S A 2005, 102, 15545–15550, doi:10.1073/pnas.0506580102.

35. Korotkevich, G.; Sukhov, V.; Budin, N.; Shpak, B.; Artyomov, M.N.; Sergushichev, A. Fast gene set enrichment analysis. bioRxiv 2021, 060012, doi:10.1101/060012.

36. Liberzon, A.; Subramanian, A.; Pinchback, R.; Thorvaldsdóttir, H.; Tamayo, P.; Mesirov, J.P. Molecular signatures database (MSigDB) 3.0. Bioinformatics 2011, 27, 1739–1740, doi:10.1093/bioinformatics/btr260.

37. Alexa, A.; Rahnenfuhrer, J. topGO: Enrichment analysis for Gene Ontology. R package version 2.28. 0. Cranio 2016, 2, 289–300.

38. Szklarczyk, D.; Gable, A.L.; Lyon, D.; Junge, A.; Wyder, S.; Huerta-Cepas, J.; Simonovic, M.; Doncheva, N.T.; Morris, J.H.; Bork, P.; et al. STRING v11: protein-protein association networks with increased coverage, supporting functional discovery in genome-wide experimental datasets. Nucleic Acids Res 2019, 47, D607–D613, doi:10.1093/nar/gky1131.

39. Cajka, T.; Fiehn, O. Increasing lipidomic coverage by selecting optimal mobile-phase modifiers in LC–MS of blood plasma. Metabolomics 2016, 12, 34, doi:10.1007/s11306-015-0929-x.

40. Bowden, J.A.; Heckert, A.; Ulmer, C.Z.; Jones, C.M.; Koelmel, J.P.; Abdullah, L.; Ahonen, L.; Alnouti, Y.; Armando, A.M.; Asara, J.M.;, et al. Harmonizing lipidomics: NIST interlaboratory comparison exercise for lipidomics using SRM 1950-Metabolites in Frozen Human Plasma. J Lipid Res 2017, 58, 2275–2288, doi:10.1194/jlr.M079012.

41. Ismail, I.T.; Showalter, M.R.; Fiehn, O. Inborn Errors of Metabolism in the Era of Untargeted Metabolomics and Lipidomics. Metabolites 2019, 9, doi:10.3390/metabo9100242.

42. Zhang, Y.; Fan, S.; Wohlgemuth, G.; Fiehn, O. Denoising Autoencoder Normalization for Large-Scale Untargeted Metabolomics by Gas Chromatography-Mass Spectrometry. Metabolites 2023, 13, doi:10.3390/metabo13080944.

43. Arce, A.; Altman, R.; Badolian, A.; Low, J.; Cuaresma, A.B.; Halkia, G.; Keshet, U.; Fiehn, O.; Stahelin, R.V.; Nikolaidis, N. Heat shock-induced PI(4)P increase drives HSPA1A translocation to the plasma membrane in cancer and stressed cells through PI4KIII alpha activation. Cell Stress Chaperones 2026, 31, 100130, doi:10.1016/j.cstres.2025.100130.

44. Low, J.; Altman, R.; Badolian, A.; Cuaresma, A.B.; Briseno, C.; Keshet, U.; Fiehn, O.; Stahelin, R.V.; Nikolaidis, N. Heat-induced phosphatidylserine changes drive HSPA1A’s plasma membrane localization. Cell Stress Chaperones 2025, 30, 100092, doi:10.1016/j.cstres.2025.100092.

45. Chong, J.; Wishart, D.S.; Xia, J. Using MetaboAnalyst 4.0 for Comprehensive and Integrative Metabolomics Data Analysis. Curr Protoc Bioinformatics 2019, 68, e86, doi:10.1002/cpbi.86.

46. Reed, A.D.; Pensa, S.; Steif, A.; Stenning, J.; Kunz, D.J.; Porter, L.J.; Hua, K.; He, P.; Twigger, A.-J.; Siu, A.J.Q.;, et al. A single-cell atlas enables mapping of homeostatic cellular shifts in the adult human breast. Nature Genetics 2024, 56, 652–662, doi:10.1038/s41588-024-01688-9.

