## Supplemental Figures S1-S4 for "Population-associated molecular variation in histologically normal breast tissue is context-dependent and associated with distinct transcriptional states"

A

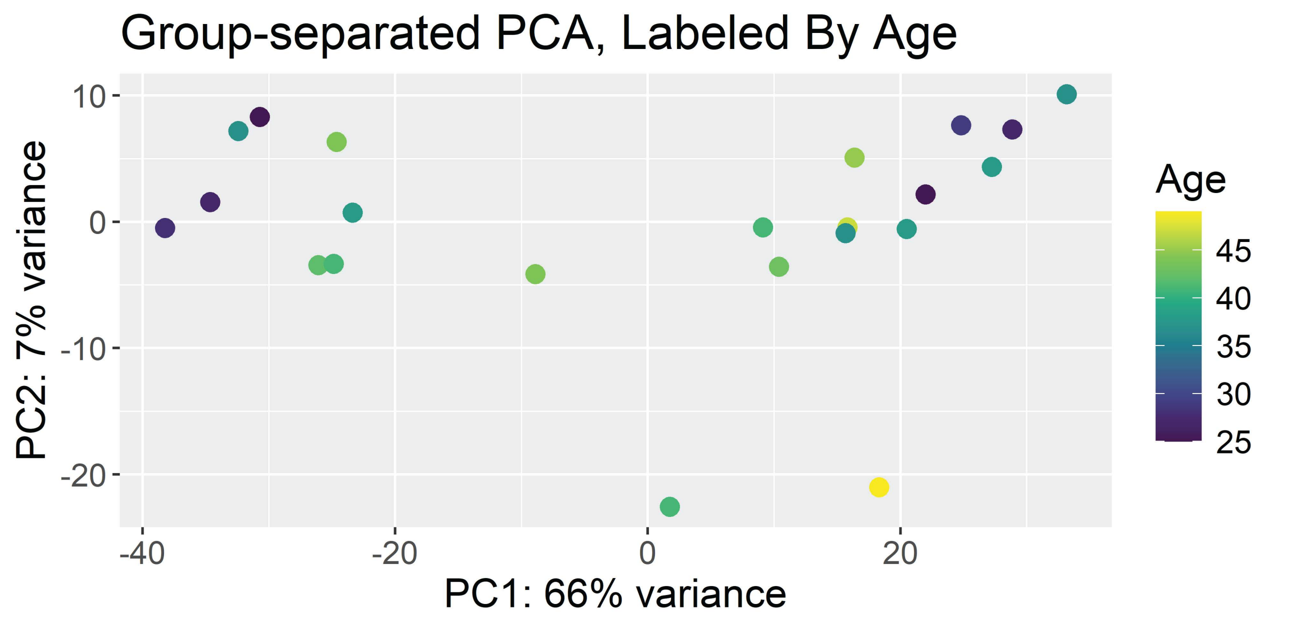
Group-separated PCA, Labeled By Age
PC2: 7% variance
PC1: 66% variance
Age
45
40
35
30
25
10
0
-10
-20
-40
-20
0
20

B

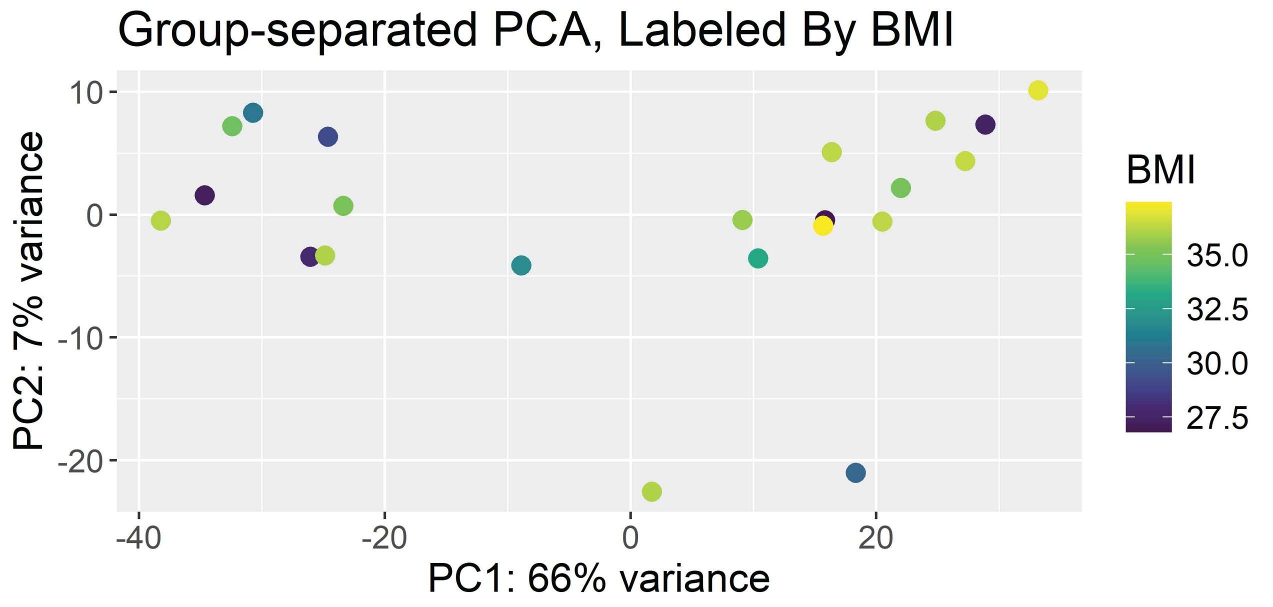
Group-separated PCA, Labeled By BMI
PC2: 7% variance
PC1: 66% variance
BMI
35.0
32.5
30.0
27.5
10
0
-10
-20
-40
-20
0
20

C

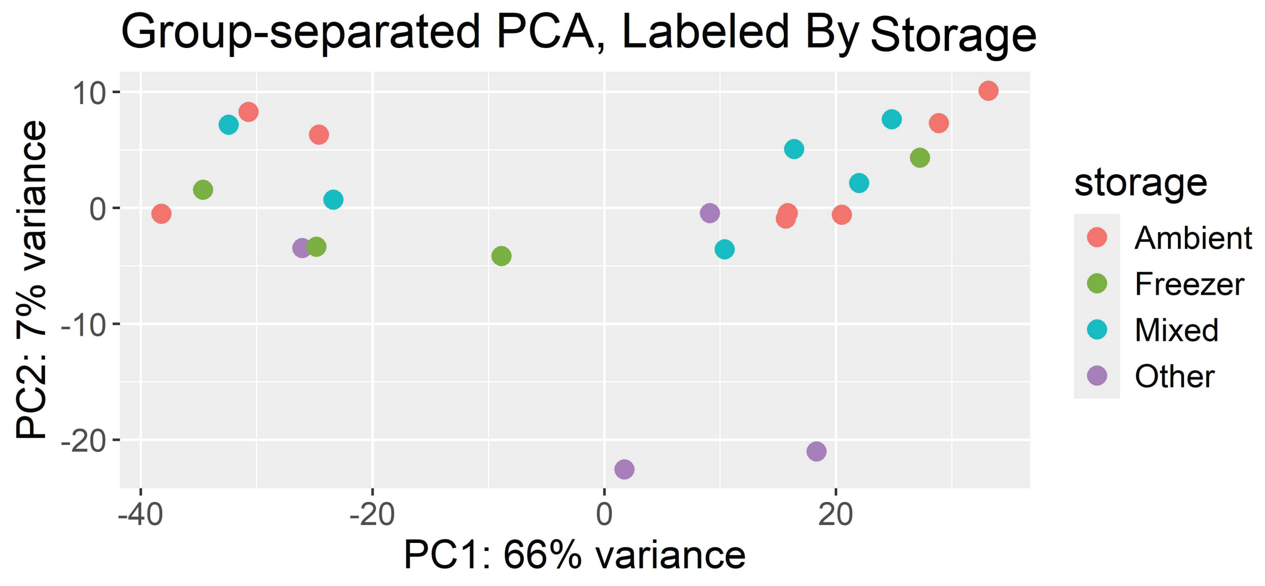
Group-separated PCA, Labeled By Storage
PC2: 7% variance
PC1: 66% variance
storage
Ambient
Freezer
Mixed
Other
10
0
-10
-20
-40
-20
0
20

**Supplementary Figure S1. Evaluation of potential metadata drivers of transcriptomic variation.** Principal component analysis (PCA) of normalized transcriptomic profiles colored by (A) age, (B) body mass index (BMI), and (C) sample storage category. Storage categories reflect sample handling conditions as recorded in metadata (e.g., ambient, freezer, mixed, other), while all samples were ultimately preserved under liquid nitrogen conditions. No distinct clustering of samples was observed with respect to these variables, indicating that the primary transcriptomic separation is not driven by these metadata factors, although subtle effects cannot be excluded given the sample size.

**A**

**Top Upregulated Pathways (Positive NES)**

| Pathway | NES | Adjusted p-value |
|---|---|---|
| GOBP_ADAPTIVE_IMMUNE_RESPONSE | 3.34 | $8.54 \times 10^{-25}$ |
| GOBP_B_CELL_MEDIATED_IMMUNITY | 3.05 | $4.82 \times 10^{-11}$ |
| GOBP_LYMPHOCYTE_MEDIATED_IMMUNITY | 2.71 | $6.13 \times 10^{-8}$ |

**Top Downregulated Pathways (Negative NES)**

| Pathway | NES | Adjusted p-value |
|---|---|---|
| GOBP_B_CELL_MEDIATED_IMMUNITY | -5.08 | 0.0375 |
| GOBP_POSITIVE_REGULATION_OF_PROTEIN _CONTAINING_COMPLEX_DISASSEMBLY | -5.20 | 0.0375 |
| GOBP_VACUOLE_ORGANIZATION | -5.29 | 0.0288 |

**B**

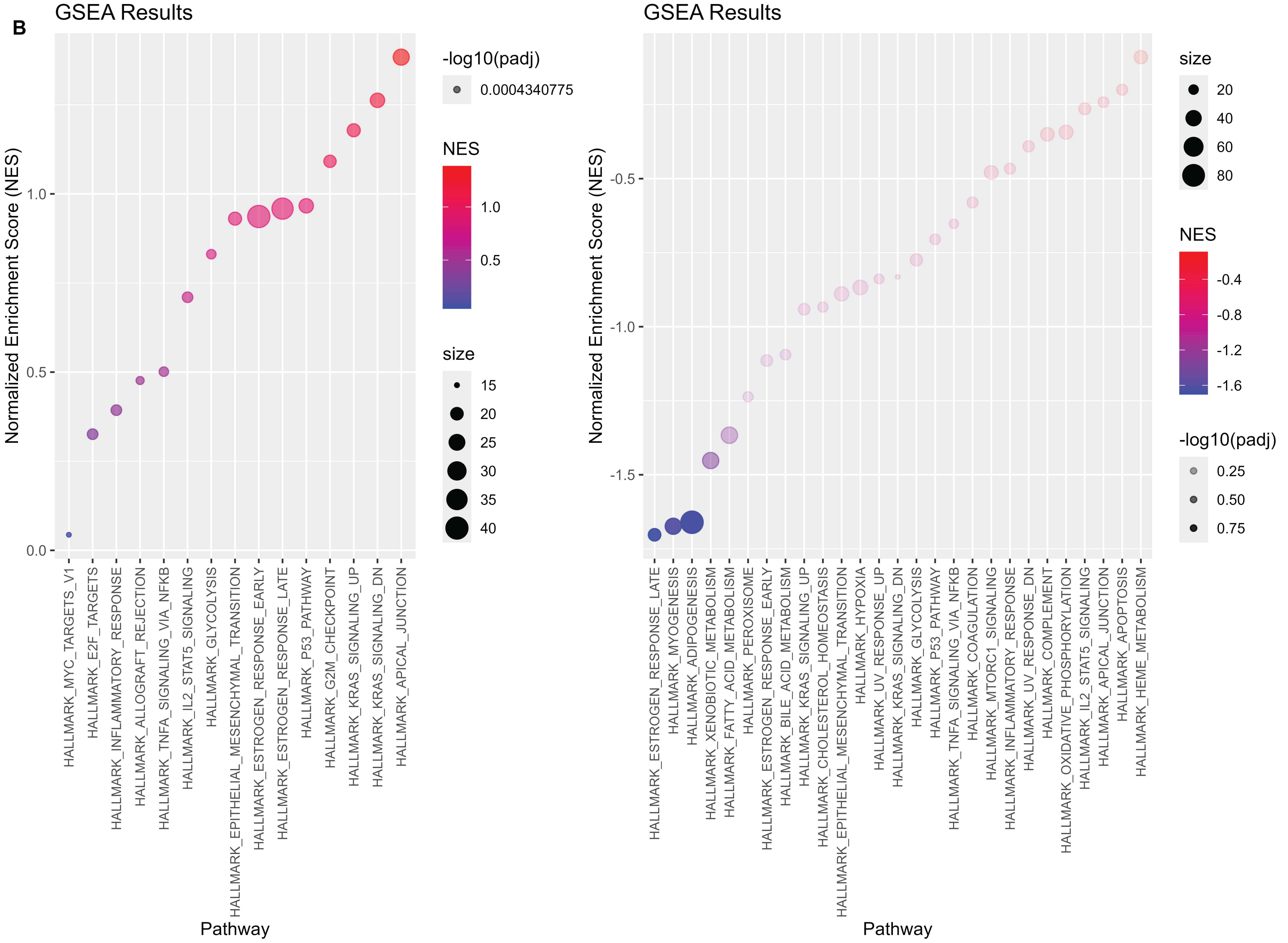


**C**

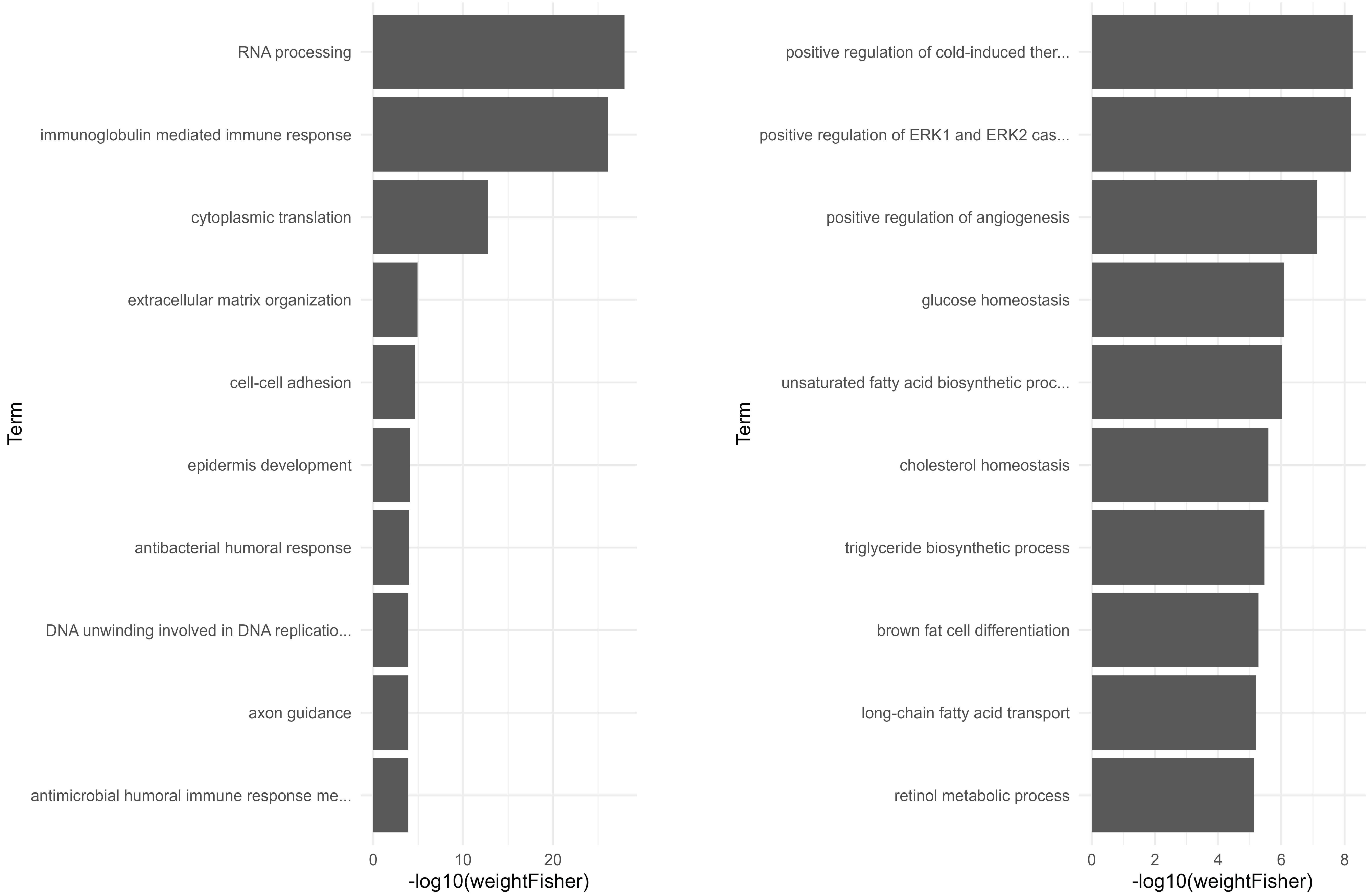

**Supplementary Figure S2. Expanded pathway enrichment analysis for transcriptomic differences between G1 and G2.** (A) Summary tables of the top significantly enriched Gene Ontology (GO) biological process pathways for positively enriched (left; higher in G1 relative to G2) and negatively enriched (right; higher in G2 relative to G1) gene sets. Pathways are ranked by normalized enrichment score (NES) and adjusted *p*-value. (B) Gene set enrichment analysis (GSEA) of Hallmark pathways showing positively enriched pathways in G1 relative to G2 (left) and negatively enriched pathways (i.e., enriched in G2; right). Points are colored by NES and sized according to gene set size, with significance indicated by −log10(adjusted *p*-value). (C) Gene Ontology (GO) enrichment analysis of differentially expressed genes using weighted Fisher's exact test. Bar plots show the top enriched biological processes for genes upregulated in G1 relative to G2 (left) and genes upregulated in G2 relative to G1 (right), ranked by −log10(weighted Fisher *p*-value).

A

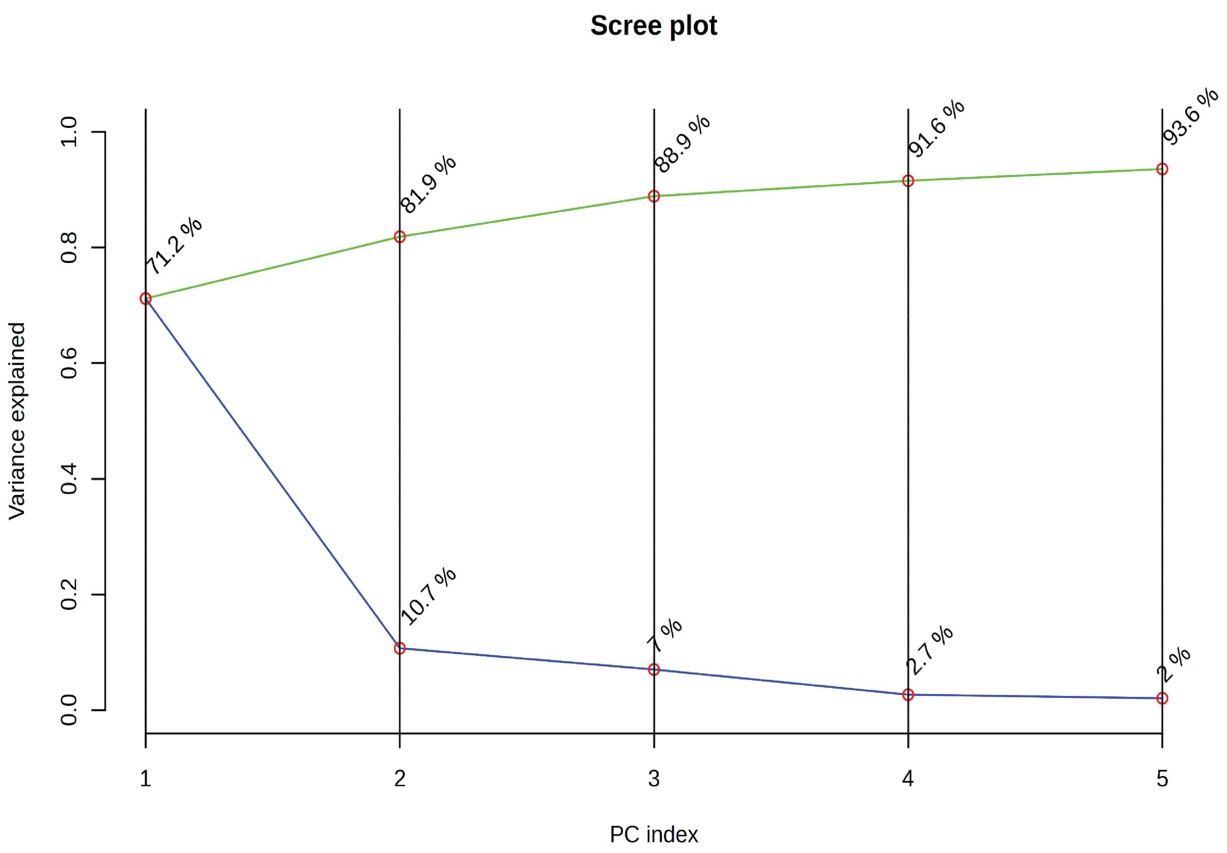

Scree plot
Variance explained
0.0
0.2
0.4
0.6
0.8
1.0
71.2 %
81.9 %
88.9 %
91.6 %
93.6 %
10.7 %
7 %
2.7 %
2 %
1
2
3
4
5
PC index


B

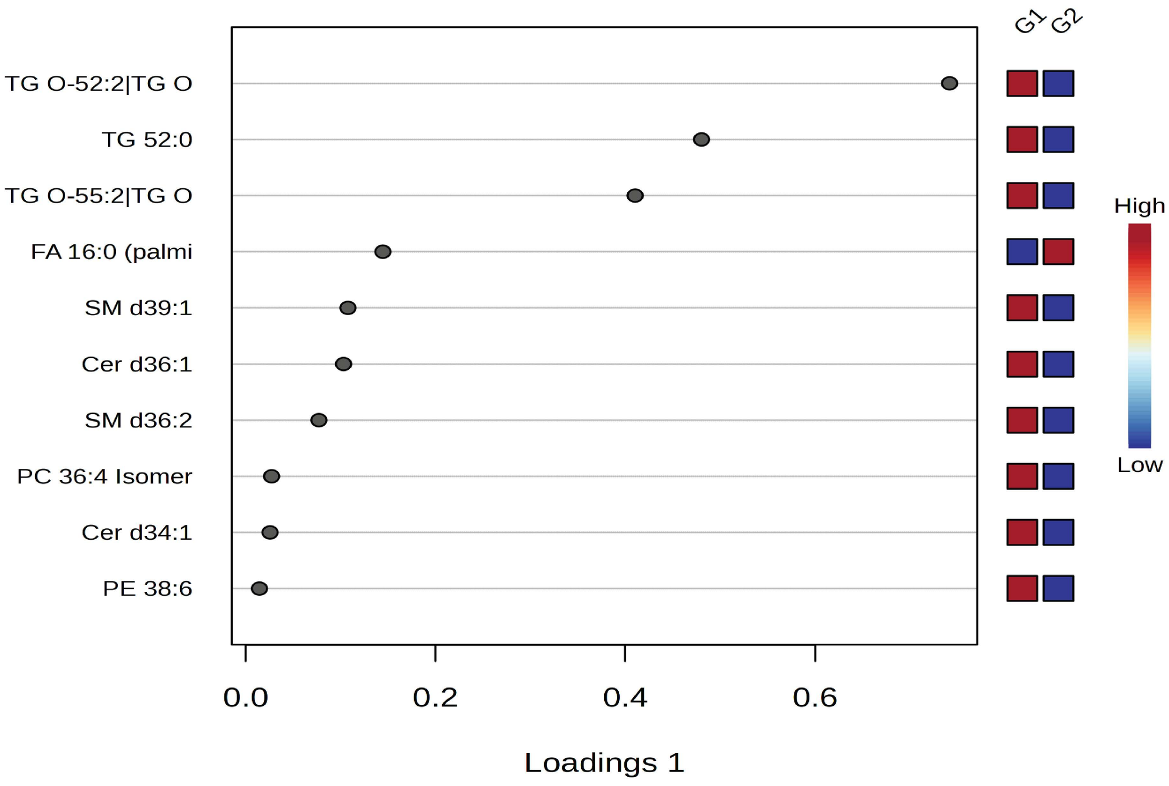

G1
G2
TG O-52:2|TG O
TG 52:0
TG O-55:2|TG O
FA 16:0 (palmi
SM d39:1
Cer d36:1
SM d36:2
PC 36:4 Isomer
Cer d34:1
PE 38:6
High
Low
0.0
0.2
0.4
0.6
Loadings 1

**Supplementary Figure S3. Principal component variance structure and lipid features contributing to group separation.** (A) Scree plot of the lipidomic principal component analysis (PCA) corresponding to Figure 4A, showing the proportion of variance explained by each principal component and the cumulative variance explained. PC1 and PC2 together account for 81.9% of the total variance. (B) Loadings from the sparse partial least squares discriminant analysis (sPLS-DA) corresponding to Figure 4B, highlighting the lipid species contributing most strongly to group separation along the first component. Relative abundance patterns for each lipid are shown for G1 and G2, with color indicating higher (red) or lower (blue) levels.

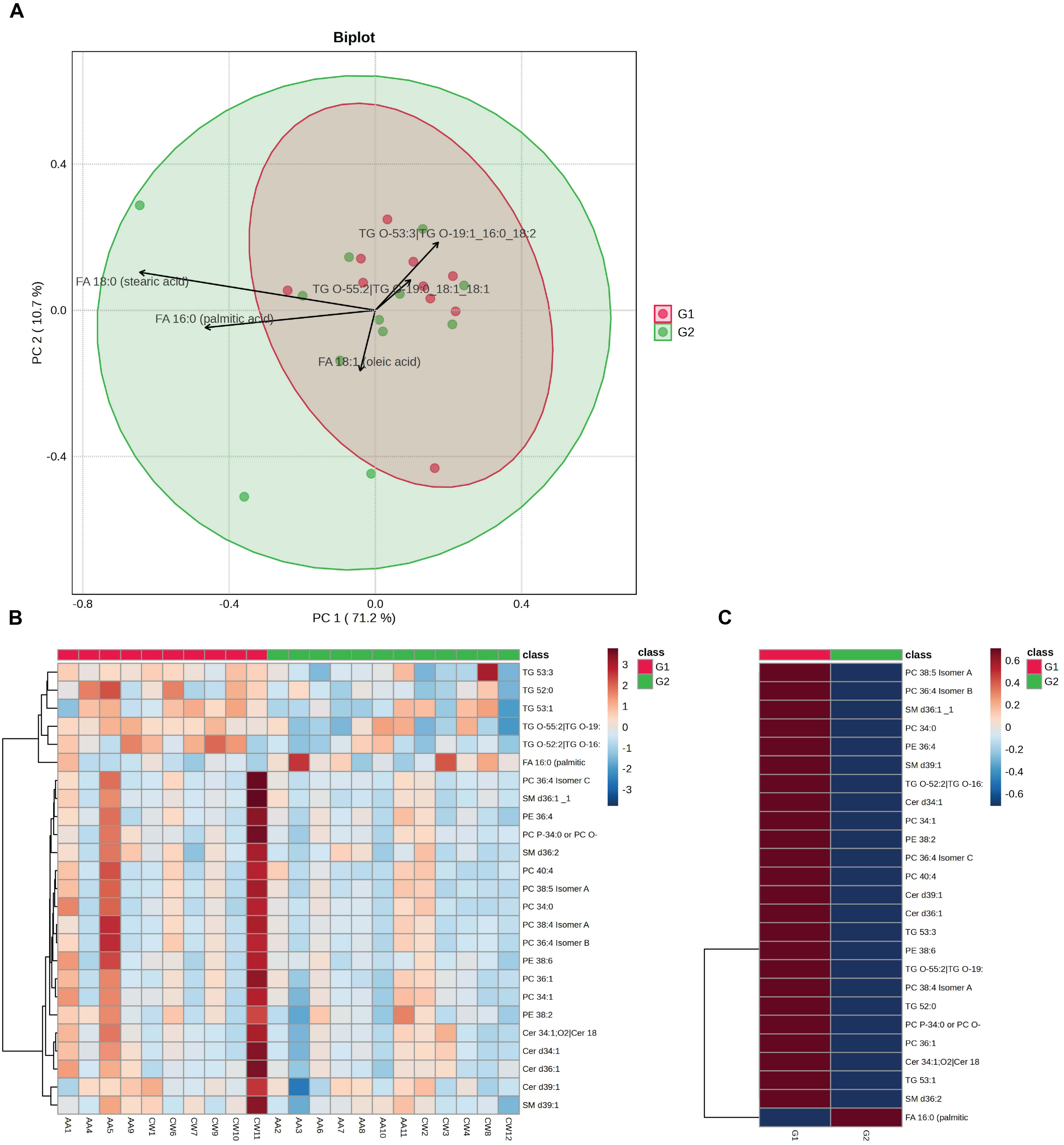
A
Biplot
PC 1 ( 71.2 %)
PC 2 ( 10.7 %)
-0.8
-0.4
0.0
0.4
TG O-53:3|TG O-19:1_16:0_18:2
FA 18:0 (stearic acid)
TG O-55:2|TG O-19:0_18:1_18:1
FA 16:0 (palmitic acid)
FA 18:1 (oleic acid)
G1
G2
B
class
G1
G2
3
2
1
0
-1
-2
-3
TG 53:3
TG 52:0
TG 53:1
TG O-55:2|TG O-19:
TG O-52:2|TG O-16:
FA 16:0 (palmitic
PC 36:4 Isomer C
SM d36:1 _1
PE 36:4
PC P-34:0 or PC O-
SM d36:2
PC 40:4
PC 38:5 Isomer A
PC 34:0
PC 38:4 Isomer A
PC 36:4 Isomer B
PE 38:6
PC 36:1
PC 34:1
PE 38:2
Cer 34:1;O2|Cer 18
Cer d34:1
Cer d36:1
Cer d39:1
SM d39:1
AA1
AA4
AA5
AA9
CW1
CW6
CW7
CW9
CW10
CW11
AA2
AA3
AA6
AA7
AA8
AA10
AA11
CW2
CW3
CW4
CW8
CW12
C
class
G1
G2
0.6
0.4
0.2
0
-0.2
-0.4
-0.6
PC 38:5 Isomer A
PC 36:4 Isomer B
SM d36:1 _1
PC 34:0
PE 36:4
SM d39:1
TG O-52:2|TG O-16:
Cer d34:1
PC 34:1
PE 38:2
PC 36:4 Isomer C
PC 40:4
Cer d39:1
Cer d36:1
TG 53:3
PE 38:6
TG O-55:2|TG O-19:
PC 38:4 Isomer A
TG 52:0
PC P-34:0 or PC O-
PC 36:1
Cer 34:1;O2|Cer 18
TG 53:1
SM d36:2
FA 16:0 (palmitic
G1
G2

**Supplementary Figure S4. Multivariate and feature-level lipidomic differences between G1 and G2.** (A) Principal component analysis (PCA) biplot of lipidomic profiles showing partial separation between G1 and G2 samples. PC1 and PC2 explain 71.2% and 10.7% of the variance, respectively. Arrows indicate lipid species contributing to group separation, including fatty acids (FA 16:0, FA 18:0, FA 18:1) and ether-linked triglycerides including TG O-53:3 and TG O-55:2. Ellipses represent group distributions. (B) Heatmap of selected lipid species across all samples, with hierarchical clustering of lipids. Values are scaled as Z-scores. Samples are annotated by group (G1 vs G2). Patterns indicate modest but consistent differences in specific lipid subsets between groups. (C) Heatmap of top discriminating lipids identified by sPLS-DA. Lipids enriched in G1 include phospholipids and sphingomyelins, while G2 is associated with triglycerides and ether-linked triglycerides. Values are shown as scaled intensities (Z-scores), illustrating directional differences despite limited statistical significance.
