## Supplemental Tables S1-S9 for "Population-associated molecular variation in histologically normal breast tissue is context-dependent and associated with distinct transcriptional states": STableS3.docx

**Supplementary Table S3.** Cell-type deconvolution of bulk RNA-seq data using MuSiC. Cell-type proportions were estimated from bulk RNA-seq data (n = 22) using MuSiC with a normal human mammary gland single-cell reference. Individual cell types were collapsed into four categories: Epithelial (luminal epithelial, basal-myoepithelial, and secretory precursor cells), Vascular (blood vessel and lymphatic endothelial cells), Stromal (fibroblasts and related mesenchymal cells), and Immune (leukocytes). Group differences were assessed using two-sided Wilcoxon rank-sum tests with Benjamini-Hochberg correction. Values are medians of estimated proportions (scale 0–1).

**Part A — G1 vs G2 comparison (n = 10 vs n = 12)**

| **Category** | **Component cell types** | **G1 median** | **G2 median** | **p-value** | **padj** | **Sig.** |
| --- | --- | --- | --- | --- | --- | --- |
| Epithelial | Luminal epithelial + basal-myoepithelial + secretory precursor | 0.802 | 0.609 | 0.0001 | 0.0003 | ****** |
| Vascular | Blood vessel endothelial + lymphatic endothelial | 0.176 | 0.368 | 0.0001 | 0.0003 | ****** |
| Stromal | Fibroblast + mural + perivascular | 0.009 | 0.043 | 0.457 | 0.610 | **ns** |
| Immune | Leukocyte | 0.000 | 0.000 | 0.867 | 0.867 | **ns** |

*G1 median > G2: epithelial-enriched baseline state. G2 median > G1: vascular-enriched baseline state.*

*** padj < 0.001	ns = not significant*

**Part B — AA vs CW within G1 (n = 4 AA vs n = 6 CW)**

**Note:** The primary purpose of this comparison is to assess whether population-associated transcriptional differences observed within G1 could be explained by differences in cellular composition. Non-significance indicates that the transcriptional signal is not attributable to compositional differences between groups.

| **Category** | **Component cell types** | **AA median** | **CW median** | **p-value** | **padj** | **Sig.** |
| --- | --- | --- | --- | --- | --- | --- |
| Epithelial | Luminal epithelial + basal-myoepithelial + secretory precursor | 0.800 | 0.802 | 0.914 | 0.914 | **ns** |
| Vascular | Blood vessel endothelial + lymphatic endothelial | 0.162 | 0.188 | 0.610 | 0.813 | **ns** |
| Stromal | Fibroblast + mural + perivascular | 0.020 | 0.007 | 0.450 | 0.813 | **ns** |
| Immune | Leukocyte | 0.000 | 0.000 | 0.359 | 0.813 | **ns** |

*No significant differences in estimated cell-type proportions between AA and CW samples within G1 (all FDR-adjusted p > 0.81), indicating that population-associated transcriptional differences are not driven by cellular composition differences.*

*ns = not significant after Benjamini-Hochberg correction*

**Part C — Original individual cell type results (informational)**

Results from Wilcoxon tests on individual cell types prior to collapsing. Four cell types returned NA (zero variance across all samples) and are excluded. These results are provided for transparency; the collapsed analysis in Parts A and B is the primary reported analysis.

**G1 vs G2**

| **Cell type** | **G1 median** | **G2 median** | **p-value** | **padj** | **Sig.** |
| --- | --- | --- | --- | --- | --- |
| Luminal epithelial cell of mammary gland | 0.624 | 0.499 | 0.023 | 0.069 | ***** |
| Basal-myoepithelial cell of mammary gland | 0.198 | 0.124 | 0.138 | 0.276 | **ns** |
| Blood vessel endothelial cell | 0.122 | 0.359 | 0.0001 | 0.0007 | ****** |
| Endothelial cell of lymphatic vessel | 0.003 | 0.001 | 0.380 | 0.549 | **ns** |
| Fibroblast of mammary gland | 0.009 | 0.043 | 0.457 | 0.549 | **ns** |
| Leukocyte | 0.000 | 0.000 | 0.867 | 0.867 | **ns** |

*** padj < 0.01	* padj < 0.10	ns = not significant*

**AA vs CW within G1**

| **Cell type** | **AA median** | **CW median** | **p-value** | **padj** | **Sig.** |
| --- | --- | --- | --- | --- | --- |
| Basal-myoepithelial cell of mammary gland | 0.228 | 0.201 | 0.610 | 0.711 | **ns** |
| Blood vessel endothelial cell | 0.145 | 0.123 | 0.610 | 0.711 | **ns** |
| Endothelial cell of lymphatic vessel | 0.002 | 0.039 | 0.914 | 0.914 | **ns** |
| Fibroblast of mammary gland | 0.034 | 0.012 | 0.450 | 0.711 | **ns** |
| Leukocyte | 0.001 | 0.000 | 0.359 | 0.711 | **ns** |
| Luminal adaptive secretory precursor | 0.001 | 0.000 | 0.094 | 0.659 | **ns** |
| Luminal epithelial cell of mammary gland | 0.632 | 0.637 | 0.610 | 0.711 | **ns** |

*ns = not significant after Benjamini-Hochberg correction. All padj > 0.65.*
